# Phazolicin – a Novel Thiazole/Oxazole-Modified Peptide Inhibiting the Bacterial Ribosome in a Species-Specific Way

**DOI:** 10.1101/705970

**Authors:** Dmitrii Y. Travin, Zoe L. Watson, Mikhail Metelev, Fred R. Ward, Ilya A. Osterman, Irina M. Khven, Nelli F. Khabibullina, Marina Serebryakova, Peter Mergaert, Yury S. Polikanov, Jamie H. D. Cate, Konstantin Severinov

**Affiliations:** Center of Life Sciences, Skolkovo Institute of Science and Technology, Moscow, Russia; Institute of Gene Biology, Russian Academy of Science, Moscow, Russia; Department of Chemistry, University of California, Berkeley, USA; Department of Molecular and Cell Biology, University of California, Berkeley, USA; Department of Chemistry, Lomonosov Moscow State University, Moscow, Russia; Department of Bioengineering and Bioinformatics, Lomonosov Moscow State University, Moscow, Russia; Department of Biological Sciences, University of Illinois at Chicago, Chicago, Illinois, USA; A.N. Belozersky Institute of Physico-Chemical Biology, Lomonosov Moscow State University, Moscow, Russia; Institute for Integrative Biology of the Cell (I2BC), CNRS CEA University Paris-Sud, University Paris-Saclay, Gif-sur-Yvette, France; Department of Medicinal Chemistry and Pharmacognosy, University of Illinois at Chicago, Chicago, USA; Waksman Institute for Microbiology, Rutgers, the State University of New Jersey, Piscataway, USA

**Keywords:** Phazolicin, azole-containing peptide, translation inhibitor, RiPPs, *Rhizobium* natural products, cryo-EM structure, ribosome

## Abstract

Ribosomally synthesized and post-translationally modified peptides (RiPPs) are a rapidly expanding and largely untapped class of natural products with various biological activities. Linear azol(in)e-containing peptides (LAPs) comprise a subclass of RiPPs that display an outstanding diversity of mechanisms of action while sharing common structural features. Here, we report the discovery of a new LAP biosynthetic gene cluster in the genome of *Rhizobium sp.* Pop5, which encodes the precursor peptide and modification machinery of phazolicin (PHZ) – an extensively modified peptide exhibiting narrow-spectrum antibacterial activity against some symbiotic bacteria of leguminous plants belonging to the *Rhizobiales*. PHZ inhibits prokaryotic translation through the obstruction of the passage of the nascent peptide through the ribosome exit channel. The cryo-EM structure of the *Escherichia coli* ribosome with bound PHZ revealed that the drug interacts with the 23S rRNA and ribosomal proteins uL4 and uL22 and obstructs the exit tunnel in a way that is distinct from other compounds blocking the exit channel. We show that the sequence of uL4 ribosomal protein loop involved in PHZ binding determines the species-specificity of antibiotic interaction with its target. PHZ and its predicted homologs from other bacterial species expand the known diversity of LAPs and may be used in the future as biocontrol agents for the needs of agriculture.

## INTRODUCTION

Since the discovery of the first antibiotics, natural products of bacterial and fungal origin serve as the predominant source of new compounds with various in-high-demand functional activities in the fight against ever-growing antibiotic resistance and cancer^1,2^. While traditional activity-based screens often lead to the rediscovery of already known compounds, the ever-increasing genomic data make genome-mining based strategies viable alternatives in the search for new bioactive compounds. Ribosomally synthesized and post-translationally modified peptides (RiPPs) comprise a diverse class of natural products that are well suited for genome-mining approaches as their biosynthetic gene clusters (BGCs) contain multiple signature genes coding for enzymes performing posttranslational modifications and the peptide precursor genes, allowing to significantly improve the quality of bioinformatics predictions^3^.

Linear azol(in)e-containing peptides (LAPs) form a subfamily of RiPPs sharing thiazol(in)e and (methyl)oxazol(in)e heterocycles, which result from cyclization of Cys and Ser (Thr) side chains during the posttranslational modification of the precursor peptide^4^. In addition to the precursor peptide-coding gene (gene A) LAP BGCs typically include genes for an YcaO-domain containing cyclodehydratase (gene D), its partner protein (gene C or gene F), FMN-dependent dehydrogenase (gene B), which oxidizes azolines to azoles, and the efflux pump^5^. Although less than two dozens of LAPs can be considered as well-characterized, they demonstrate antibacterial activity through several completely different mechanisms. For example, microcin B17 is a DNA-gyrase poison^6^, plantazolicin targets the membrane^7^, and klebsazolicin inhibits the ribosome^8^. This diversity of modes of action, unprecedented for peptides sharing common chemical features, makes LAPs a group of special interest for the search of new antibacterials.

Here, we report the discovery of a new LAP BGC in the genome of *Rhizobium sp.* Pop5, and characterization of its product, phazolicin (PHZ) – a novel linear thiazole/oxazole-modified peptide, which targets the bacterial ribosome. The producing strain, a symbiotic nitrogen-fixing bacterium was isolated in a tropical forest in Los Tuxtlas (Mexico) from the root nodules of wild beans (*Phaseolus vulgaris*, hence “phazolicin”). Although the studies of natural products with antibacterial properties produced by rhizobia date back to the 1960s^9^, the number of well-characterized compounds is still very small with trifolitoxin produced by *R. leguminosarum* bv. *trifolii* T24 being the only rhizobial thiazole/oxazole-containing RiPP described to date^10^. Being active against a variety of bacteria from *Rhizobiales* including several phytopathogens such as *Agrobacterium tumifaciens* and *A. rhizogenes*, phazolicin is not only interesting because of its idiosyncratic mode of interaction with the ribosome, but also as a potential agent for biocontrol in agriculture.

## RESULTS

### Identification of a new LAP biosynthetic gene cluster

Klebsazolicin (KLB) from *Klebsiella pneumoniae* is a recently characterized LAP, which inhibits prokaryotic translation through the obstruction of the ribosome exit tunnel^8^. The BGC of this RiPP (*klpACBDE*) contains genes encoding the precursor peptide (*klpA*), the three enzymes involved in the formation of azole cycles (*klpC*, *klpB,* and *klpD*) as well as a transmembrane pump (*klpE*) exporting the modified peptide. We used the amino acid sequences of KlpB and KlpD proteins as queries to search the NCBI non-redundant protein database using the PSI-BLASTP algorithm^11^ followed by manual analysis of the genomic environment of the identified gene orthologues. Along with several previously described clusters of KLB homologs^8^ a distinct BGC *phzEACBD* was found in the genome of *Rhizobium* sp. Pop5. Two other related BGCs were found in the genomes of *Rhizobium sp.* PDO1-076 and *Phyllobacterium myrsinacearum* DSM 5893, also belonging to the order *Rhizobiales*. The alignment of amino acid sequences of the predicted core parts of precursor peptides revealed a similar pattern of Cys/Ser/Thr residues, which potentially could be cyclized in the mature LAP product (**Figure 1B**, red). The putative PhzA precursor peptide has an amino acid sequence significantly different from that of the KLB precursor KlpA (**Figure 1A**). As is the case with KlpA, the C-terminal part of the predicted PhzA precursor peptide is enriched in Ser, Thr, and Cys residues but also contains several positively charged residues absent in the core part of KlpA. Moreover, PhzA and its relatives lack the X-X-(S/T)P motif required for the formation of the N-terminal amidine cycle, which is strictly required for the activity of KLB^12^. Therefore, we hypothesized that the *phzEACBD* and similar BGCs from other *Rhizobiales* might encode closely related LAPs with structural and functional properties different from those of KLB or other known compounds of the family.

**Figure 1.**
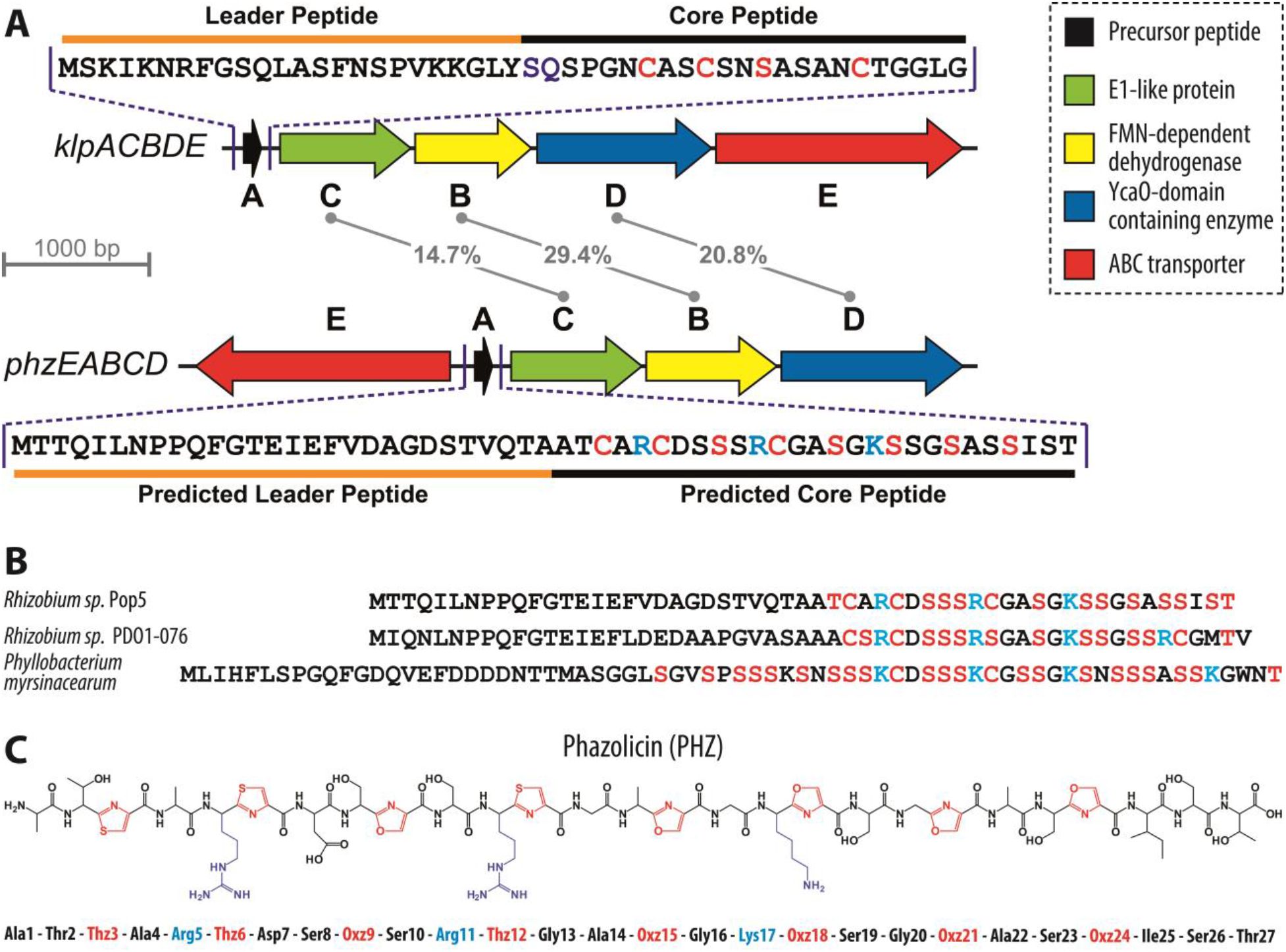
Organization of the phazolicin (PHZ) biosynthetic gene cluster. (**A**) Comparison of klebsazolicin (KLB) biosynthetic gene cluster (*klpACBDE*) with the cluster found in the genome of *Rhizobium sp.* Pop5. Functions of the genes forming the BGCs are listed on the right. The numbers in the middle indicate the extent of identity between the amino acid sequences of the C, B, and D gene products. The sequence of the KLB precursor peptide is shown above its BGC with the leader and core parts indicated. Residues converted to azoles in the final product are highlighted in red. Residues involved in the formation of the N-terminal amidine cycle are highlighted in violet. The sequence of the precursor peptide from the newly identified BGC is shown below it. Ser and Cys residues in the sequence of PhzA precursor involved in oxazole and thiazole formation respectively are shown in red. Positively charged residues of the predicted core part are highlighted in blue. (**B**) Alignment of the amino acid sequences of precursor peptides encoded in *phzEACBD* BGC and BGCs of PHZ homologs from other *Rhizobiales* (*Rhizobium sp.* PDO1-076 and *Phyllobacterium myrsinacearum* DSM 5893). Ser, Thr and Cys residues in the predicted core parts of the peptides are shown in red. Positively charged residues are highlighted in blue. (**C**) Structure of the fully modified PHZ peptide after the cleavage of the leader. Cyclized residues are shown in red (Thz and Oxz). Positively charged amino acids are highlighted in blue.

### Purification and characterization of a new LAP

Reverse-phase HPLC analysis of medium after cultivation of the *Rhizobium sp.* Pop5 led to the identification of several fractions absorbing at 254 nm, a wavelength previously used in the purification of other azole-containing compounds (**Figure S1A**)^8,13^. MALDI-TOF-MS analysis of the selected fractions showed the presence of compounds, which, based on the observed masses, appeared to be the products of the PhzA precursor peptide posttranslational modification (**Figure S1B**). The major compound with monoisotopic MH^+^ 2363.9 was named phazolicin (PHZ) (**Figure S1C**). This mass matches the mass of the C-terminal part of the PhzA precursor peptide, if it was cleaved between the two alanine residues […Thr-Ala-↓-Ala-Thr-Cys…] and contained eight azole rings (loss of 20 Da per installation of each cycle corresponds to −18 Da for each dehydration/cyclization reaction and −2 Da for each azoline oxidation event) (**Figure 1C**). Two other major compounds detected in the growth media **(Figure S1B)** also contained eight azole cycles but had an alternative site for the leader peptide cleavage situated one or two residues closer to the N-terminus of the precursor. These compounds are further referred as A-PHZ (MH^+^=2434.9) and TA-PHZ (MH^+^=2536.0). High-resolution MS measurements for PHZ, A-PHZ, and TA-PHZ resulted in MW values of 2362.8679, 2433.9052 and 2534.9524, which are all within 1.5 ppm of the corresponding masses calculated based on the brutto-formulas **(Figure S2)**. Although due to extensive modification the fragmentation of the PHZ was relatively poor, MS/MS analysis using both ESI-MS/MS and MALDI-MS/MS techniques allowed predictions of the positions of azole cycles in the structure of phazolicin **(Figure S3)**. PHZ is a 27-residue long peptide with three thiazole and five oxazole cycles in its structure. The cycles are distributed evenly across the core part of the precursor peptide with every third residue being turned into an azole **(Figure 1C)**. In addition to the major fractions, several minor forms with a fewer (5-7) number of azole cycles were detected in the cultivation medium **(Figure S1B)**. Compounds with less than eight cycles lacked the C-terminal cycles, consistent with the N- to C-terminus direction of modification of the PHZ precursor peptide.

### PHZ demonstrates antimicrobial activity against a variety of Rhizobiales

When the purified phazolicin was tested against *Escherichia coli* BW25113 grown on rich solid medium, small growth inhibition zones were observed only when the compound was applied in high concentrations (5-10 mM). Many known RiPPs demonstrate a narrow spectrum of antibacterial activity and are usually active against bacteria, which are evolutionary close to the producing strain^7,14,15^ serving as powerful weapons in the competition for the ecological niche between closely related species. Accordingly, we tested PHZ against a panel of microorganisms from the order *Rhizobiales* (class *Alphaproteobacteria*) grown in a rich liquid medium. The compound proved to be highly active against bacteria from genera *Rhizobium*, *Sinorhizobium (Ensifer)*, and *Azorhizobium* with the minimal inhibitory concentrations (MICs) being around 1 μM (**Table 1**). Phazolicin is less active against two tested strains of *Agrobacterium*, while microorganisms from the genus *Mesorhizobium* appeared to be PHZ-resistant (**Table 1**). PHZ was also tested against a small set of plant pathogens, plant-associated and soil microorganisms from *Gammaproteobacteria (Erwinia amylovora, Pantoea ananatis* PA4*, Pseudomonas putida* KT2440*, P. fluorescens* Pf-5), *Firmicutes (Bacillus subtilis* 168*, B. cereus* ATCC 4342), and *Actinobacteria* (*Arthrobacter* sp. ATCC21022, *Microtetraspora glauca* NRRL B-3735). However, no activity was observed. Taken together, these observations demonstrate that, similarly to a number of other LAPs, phazolicin is a narrow-spectrum antimicrobial compound specifically targeting several genera from *Rhizobiales*.

**Table 1.**
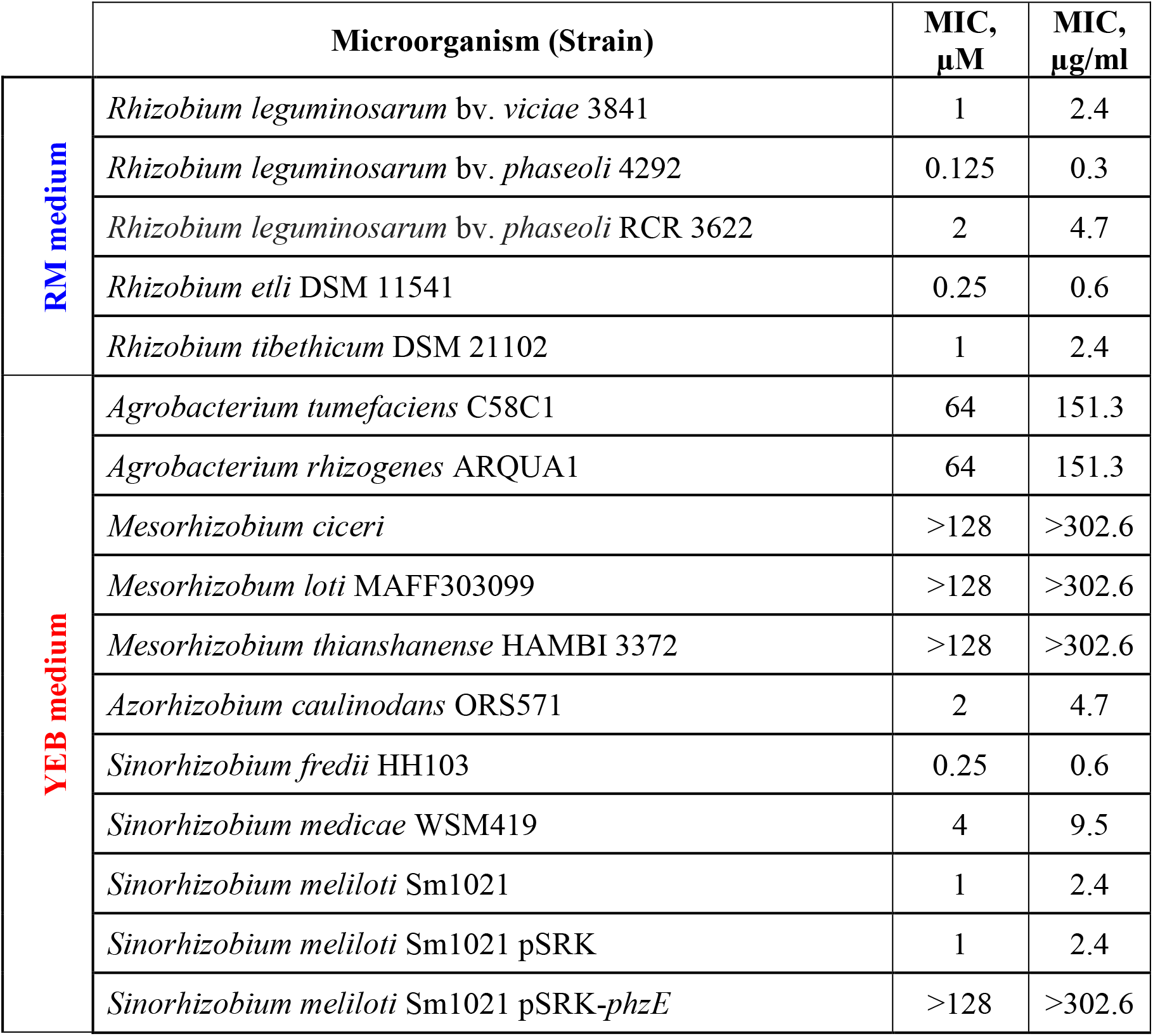
Phazolicin MICs for various bacterial strains in rich YEB medium and *Rhizobium* medium (RM).

The resistance of producing strains to RiPPs with antimicrobial activity often results from the efficient export of the mature compound by a specific transporter encoded in the corresponding BGC^16^. However, there are examples of BGCs containing additional genes providing self-immunity through different mechanisms^17,18^. Expression of a plasmid-borne copy of the *phzE* gene, which was annotated as an exporter pump, in *S. meliloti* Sm1021 conferred resistance to external PHZ and increased the MIC more than 100-fold in comparison with the strain carrying an empty vector (**Table 1**). These data confirm the role of PhzE transporter in self-immunity of the producing strain to phazolicin.

### PHZ inhibits bacterial protein translation both in vivo and in vitro

To determine the mode of action of phazolicin we tested its activity *in vivo* using an *E. coli*-based double-reporter system that allows one to identify antibiotics inducing ribosome stalling or inhibiting DNA replication^19^. In this system, pDualrep2 plasmid harbors two genes encoding fluorescent proteins under different regulation: the red fluorescent protein (RFP) is expressed in the presence of SOS-response inducing agents (e.g., fluoroquinolones or microcin B17), while the gene for far-red protein Katushka2S is transcribed only when translation inhibitors (e.g., macrolides) are added into the medium in sublethal concentrations. Similar to KLB and erythromycin (ERY), PHZ induces expression of Katushka2S but not RFP, indicating that PHZ is an inhibitor of translation (**Figure 2A**, left panel). In this assay, we observed a difference in the size of PHZ-induced inhibition zones between the *E. coli* strain with the deleted *tolC* gene and the wild-type strain (**Figure 2A**, right panel). TolC is a major outer membrane multidrug efflux porin that is responsible for the export of various compounds from the cell including siderophores (e.g., enterobactin^20^), macrolide antibiotics^21^, and some RiPPs (e.g., microcin J25^22^). In contrast to the situation observed with PHZ, the KLB-induced inhibition zones were comparable in size between the two *tolC*^−^ and wild-type *E. coli* strains, suggesting that PHZ but not KLB is subject to TolC-dependent efflux.

**Figure 2.**
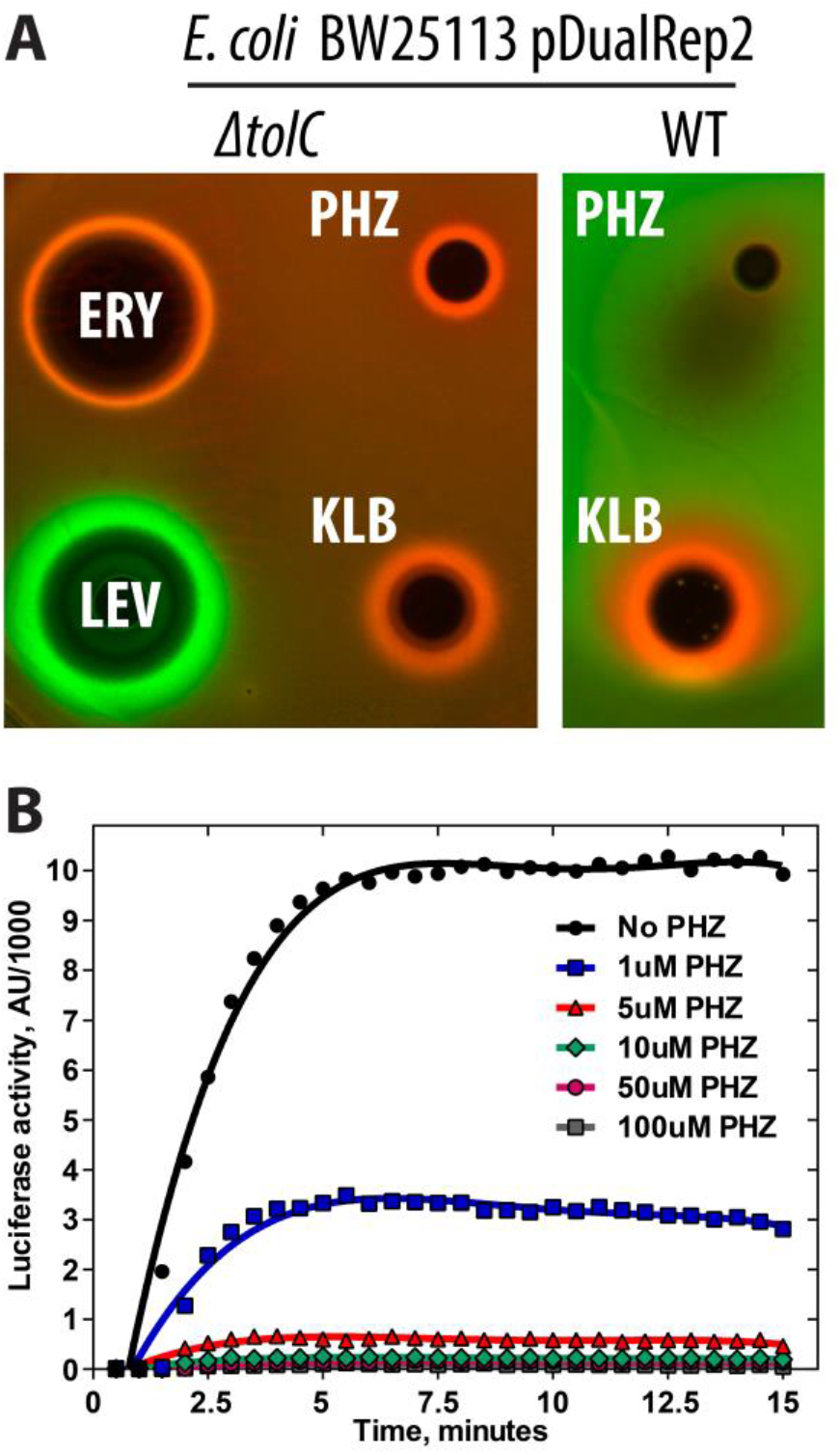
Phazolicin is an inhibitor of protein synthesis both *in vivo* and *in vitro*. (**A**) *In vivo* testing of PHZ activity using BW25113 Δ*tolC* pDualRep2 and BW25113 pDualRep2 (WT) reporter strains. The induction of RFP expression (green halo around the inhibition zone, pseudocolors) is triggered by DNA-damage while the induction of Katushka2S protein (red halo) occurs in response to ribosome stalling. Erythromycin (ERY), levofloxacin (LEV) and klebsazolicin (KLB) are used as controls. (**B**) Kinetic curves showing inhibition of protein synthesis by increasing concentrations of PHZ in the *in vitro* cell-free translation in *E. coli* S30 extract. AU, arbitrary units.

To further confirm that PHZ is a protein synthesis inhibitor, we tested its ability to decrease the translation of the luciferase mRNA *in vitro* using an *E. coli* S30 extract. Concentration-dependent inhibition of *in vitro* translation by PHZ was observed (**Figure 2B**) with 1 μM of PHZ decreasing the luciferase signal by approximately three-fold (**Figure 2B**, blue curve), and 5 μM inhibiting translation by ~95% (**Figure 2B**, red curve).

### PHZ obstructs the nascent peptide exit tunnel of the ribosome

Protein synthesis inhibitors can interfere with the activity of the ribosome or any of the various other enzymes associated with protein production (translation factors, aminoacyl-tRNA synthetases, etc). Following the analogy with KLB, we proposed that PHZ acts upon the bacterial ribosome. To determine the exact mechanism of translation inhibition by PHZ we attempted to crystallize the 70S ribosome from *Thermus thermophilus (Tth)* in complex with PHZ, mRNA, and tRNAs, as previously done with KLB^8^, and several other translation inhibitors^23–30^. Despite numerous attempts, we were not able to observe any positive electron density peaks corresponding to the ribosome-bound PHZ anywhere within the *Tth* 70S ribosome. Moreover, *in vitro* translation inhibition experiments using *E. coli (Eco)* 70S or hybrid 70S (*Eco* 30S subunit + *Tth* 50S subunit) demonstrated that PHZ does not target *Tth* ribosome (**Figure S4**). However, the same data also suggested that PHZ does target the large ribosomal subunit of the *E. coli* ribosome.

Since the crystallization-based approach was unsuccessful, we switched to cryo-EM to determine the structure of PHZ bound to the sensitive *Eco* 70S ribosome. The obtained charge density map, characterized by the overall resolution of 2.87 Å according to the “gold-standard” Fourier shell correlation (FSC) method (**Figure S5A**), revealed PHZ bound to the ribosome (**Figure 3A**). The high resolution and excellent quality of the map in the PHZ binding site (**Figure S5B**), allowed us not only to fit its atomic model (residues 2-23; **Figure 3B**) but also to confirm the positions of seven azoles in the structure of the modified peptide proposed according to the MS-MS analysis. While the results of tandem mass spectrometry indicate another oxazole present at the position 24, we left Ser23 unmodified in the model since Ser24 was not visible in the map due to its poor quality in that region (**Figure 3A**).

**Figure 3.**
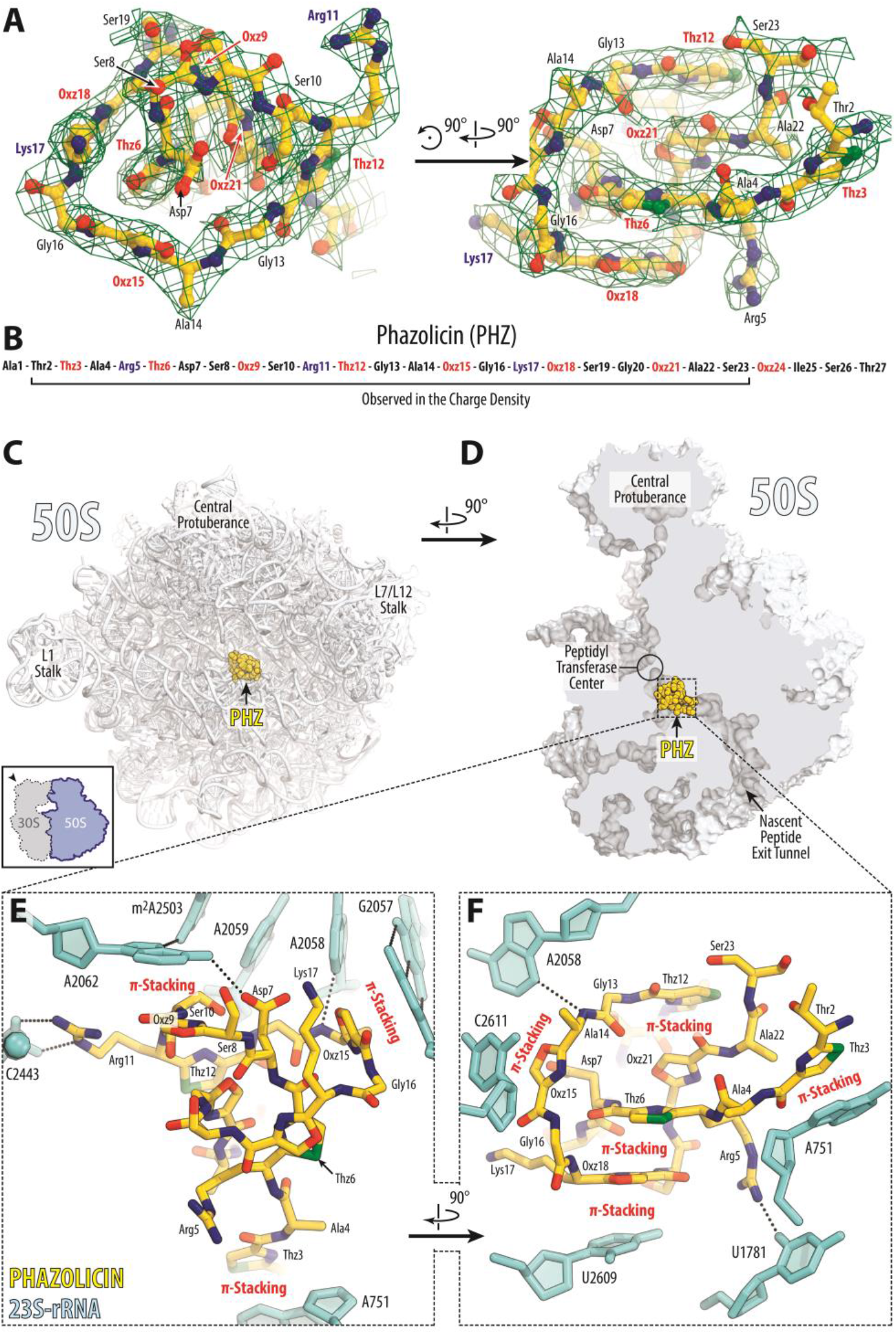
The structure of PHZ in complex with the bacterial ribosome. (**A**) Cryo-EM map of PHZ in complex with the *E. coli* 70S ribosome (green mesh). The fitted model of the compound is displayed in its respective charge density viewed from two different perspectives. The map is contoured at 2.5σ. Carbon atoms are colored yellow, nitrogen atoms are blue, oxygen atoms are red, and sulphur atoms are green. (**B**) Schematic diagram showing that only residues 2-23 of the ribosome-bound PHZ molecule are visible in the cryo-EM map. (**C, D**) Overview of the PHZ binding site (yellow) on the *E. coli* large ribosomal subunit (light blue) viewed from two different perspectives. In (C), the 50S subunit is viewed from the inter-subunit interface (30S subunit is removed for clarity) as indicated by the inset. The view in (D) is from the cytoplasm onto the A site. (**E, F**) Close-up views of the PHZ binding site in the ribosome exit tunnel. *E. coli* numbering of the nucleotides in the 23S rRNA is used. Potential H-bond interactions are indicated with dotted lines.

Similar to KLB, linear modified PHZ peptide forms a compact globule that binds in the upper part of the nascent peptide exit tunnel (NPET) of the 50S ribosomal subunit where it extensively interacts with the 23S rRNA and ribosomal proteins (**Figure 3C, D; Movie S1**). In comparison to antibiotics of smaller molecular weight such as ERY, which binds at the same location but only partially occludes the NPET (**Figure S6A, B**), binding of both KLB and PHZ results in nearly complete obstruction of the NPET (**Figure S6C, D**). A prominent feature of the ribosome-bound PHZ molecule is its complex intramolecular interactions that involves both face-to-face and edge-to-face π-π stacking of Thz12, Oxz21, Thz6, and Oxz18, along with the nucleobase U2609 from the 23S rRNA (**Figure 3F; Figure S7D**). Moreover, we observed two major types of interactions between the PHZ and the nucleotides of the 23S rRNA. First, three azoles (Thz3, Oxz15, and Oxz18) are involved in π-π stacking with nucleotides A751, C2611, and U2609, respectively (**Figure 3E, F**). Second, two positively charged side chains of Arg5 and Arg11, as well as hydrophilic side chains of Asp7 and Ser8, form additional stabilizing hydrogen bonds (H-bonds) with nucleobases and backbone phosphate groups of the 23S RNA (**Figure 3E, F; Figure S7A-C**). The presence of intramolecular π-π stacking and positively charged side chains, as well as the absence of the N-terminal amidine cycle, make the mode of PHZ binding significantly different from that of KLB. Unlike KLB, the N-terminal part of PHZ is not critical for binding to the ribosome, which is further supported by the observation that naturally occurring longer forms with an alternative leader cleavage site (A-PHZ and TA-PHZ) demonstrate the same level of *in vitro* translation inhibition activity as PHZ (**Figure S8**).

Interestingly, the PHZ binding site overlaps with those of several other classes of antibiotics such as macrolides and type B streptogramins. Comparison of the structures of ribosome-bound PHZ and KLB reveals that phazolicin binds further away than KLB from the peptidyl transferase center (PTC) of the ribosome. We superimposed our new structure of the ribosome-bound PHZ (**Figure 4A**) or the previously published KLB (**Figure 4B**) with the cryo-EM structure of the 70S ribosome containing ErmBL nascent peptide chain connected to the P-site tRNA^31^. Apparently, there is more space between the ribosome-bound drug and the PTC in the case of PHZ in comparison to KLB. This suggests that PHZ should, in principle, allow for a few more amino acids to be incorporated into the nascent polypeptide chain before it encounters the NPET-bound PHZ molecule and translation stalls. To test this hypothesis, we performed a toe-printing (primer-extension inhibition) assay, which allows identification of drug-induced ribosome stalling on a given mRNA template with a single-nucleotide precision^32^. Because PHZ causes strong induction (stalling of the ribosomes) on Dualrep2 reporter used in our *in vivo* bioactivity test (**Figure 2A**), we have chosen the corresponding mRNA (*trpL*-2Ala) as a template for our toe-printing assay. Addition of PHZ to the PURExpress cell-free transcription-translation system programmed with *trpL*-2Ala mRNA resulted in dose-dependent ribosome stalling at the seventh codon of mRNA (**Figure 4C**, lanes 4-7) while the addition of KLB induced major stalling at the third and the fifth codons (**Figure 4C**, lane 3) corroborating our hypothesis. Moreover, these data indicate that PHZ is an inhibitor of the elongation step during protein synthesis, as is the KLB. Importantly, in the presence of any of the tested antibiotics, ribosome cannot rich codons 9-10, at which it stalls in the absence of any inhibitors due to the mRNA secondary structure (**Figure 4C**, lane 8).

**Figure 4.**
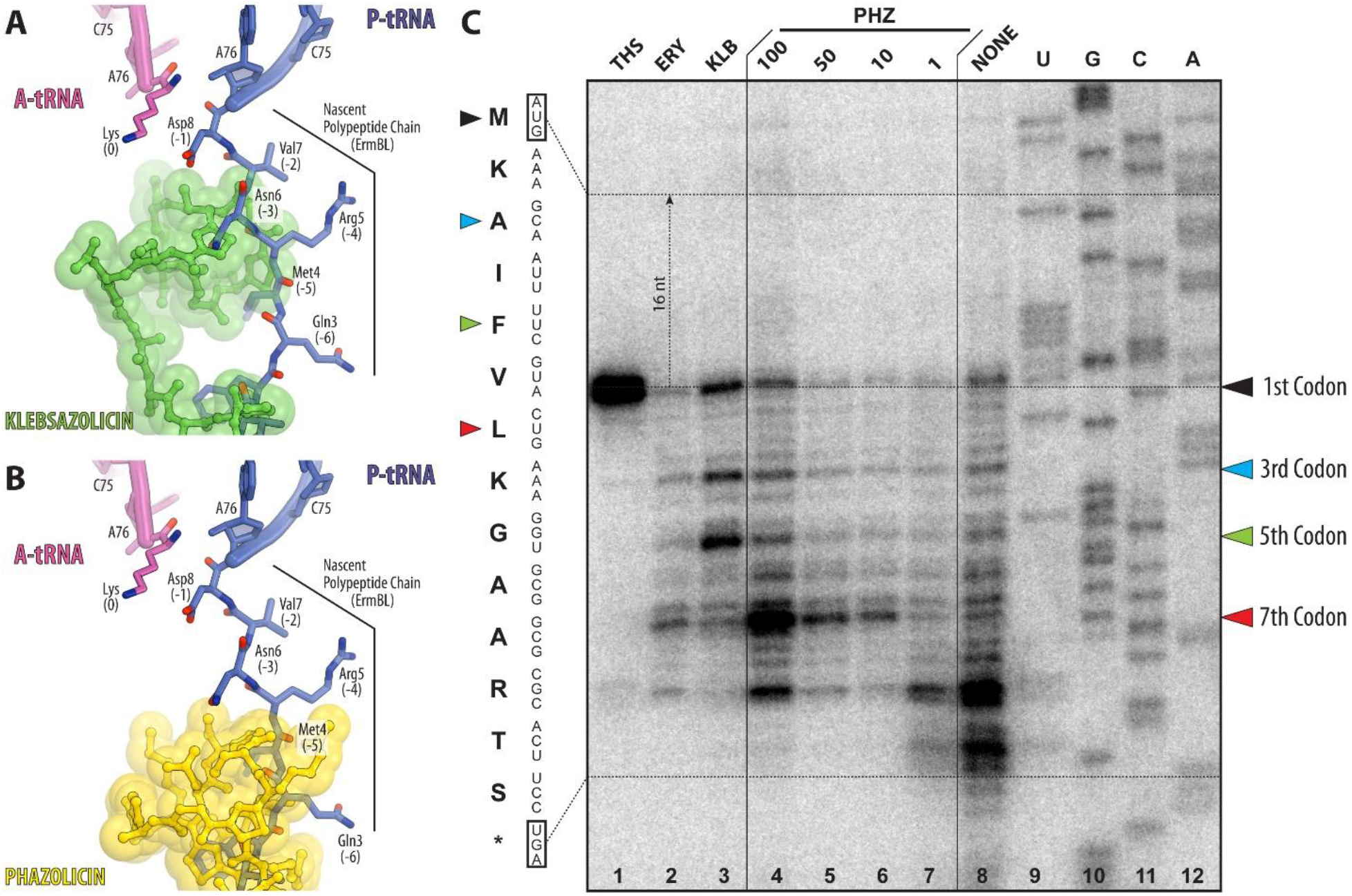
PHZ inhibits the elongation step during protein synthesis. (**A, B**) *In silico* modeling of the nascent polypeptide chain in the ribosome exit tunnel in the presence of klebsazolicin (A, PDB entry 5W4K8) or phazolicin (B, current structure) using the cryo-EM structure of the 70S ribosome with ErmBL peptide bound to the P-site tRNA (PDB entry 5JTE31). (**C**) Ribosome stalling by PHZ on *trpL-* 2Ala mRNA in comparison with other translation inhibitors (thiostrepton, THS; erythromycin, ERY; klebsazolicin, KLB), as revealed by reverse-transcription primer-extension inhibition (toe-printing) assay in a cell-free translation system. *trpL-*2Ala mRNA nucleotide sequence and the corresponding amino acid sequence are shown on the left. Colored triangles show ribosome stalling at 1^st^, 3^rd^, 5^th^ and 7^th^ codons. Note that owing to the large size of the ribosome, the reverse transcriptase stops at the nucleotide +16 relative to the codon located in the P-site.

### Amino acid sequence of the ribosomal protein uL4 loop determines species-specificity of PHZ action

The PHZ binding site is adjacent to the NPET constriction point – the most narrow part of the NPET formed by the extended loops of the ribosomal proteins uL4 and uL22 (**Figure 5A, B**). The N- and C-terminal parts of the PHZ molecule are located between the uL4 and uL22 loops (**Figure 5C**) and occupy more space in this part of the NPET than the C-terminal part of KLB (**Figure 5D**). The amino acid sequences of uL4 and uL22 proteins including their loop regions vary between distinct bacteria (**Figure 5E**). Superpositioning of our *Eco* 70S ribosome structure with bound PHZ with the structure of *Tth* 70S reveals that residues His69 of the *Tth* protein uL4 and residue Arg90 of the *Tth* protein uL22 sterically clash with the PHZ molecule (**Figure 5C**) rationalizing why PHZ does not bind to or act upon the *Tth* ribosome. Because the same residues in *Eco* ribosome are represented by less bulky Gly64 (uL4), we hypothesized that the differences in the uL4 loop sequence are responsible for the inability of PHZ to bind in the NPET and, hence, to inhibit the translation by *Tth* ribosome. To test our hypothesis, we expressed a plasmid-borne copy of the *S. meliloti rplD* gene encoding for the uL4 protein carrying the *Tth*-like G68H substitution (*rplD*^G68H^) and showed that the loop structure of *Tth* uL4 confers resistance to PHZ and allows growth of otherwise sensitive strain on the medium with 10xMIC of the antibiotic (**Figure 5F**). The expression of a double mutant *rplD*^G68H+K65A^ gene, which carries an additional substitution of the *Eco* Arg61-equivalent that forms H-bond with the PHZ in our structure, provides approximately the same level of resistance. Thus, the fine structure of the PHZ binding site (formed in part by the loop regions of ribosomal proteins uL4 and uL22) together with the repertoire of peptide transporters involved in the uptake contribute to the specificity of the antibiotic action.

**Figure 5.**
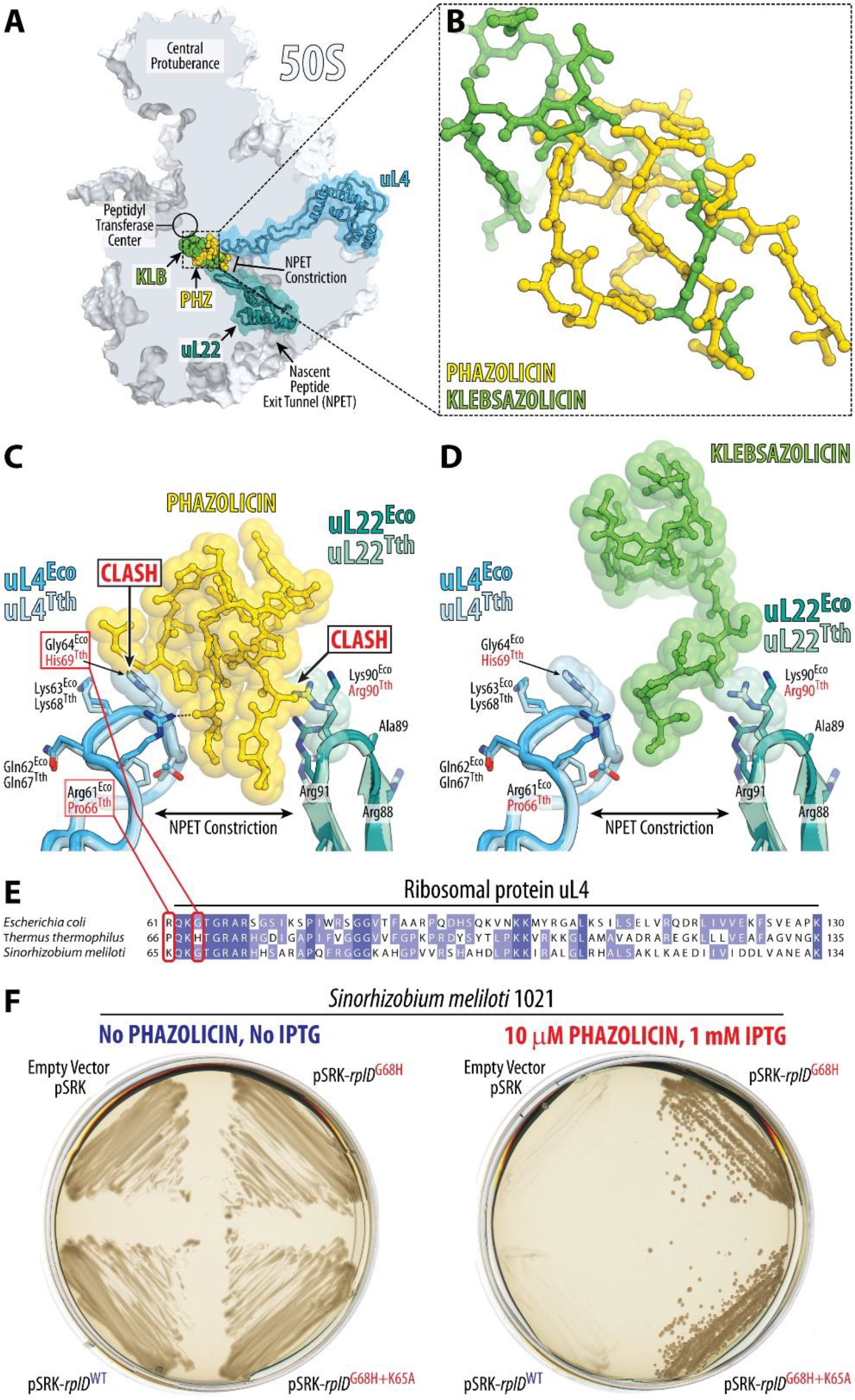
The loop of ribosomal protein uL4 defines species-specificity of PHZ action. (**A**) Comparison of PHZ (yellow) and KLB (green) binding sites in the ribosome exit tunnel. The 50S subunit is shown in light blue. Ribosomal proteins uL4 and uL22 are highlighted in blue and teal, respectively. (**B**) Superposition of the ribosome structures with bound KLB (PDB entry 5W4K^8^) and PHZ based on the alignment of the 23S rRNAs. Ribosome parts are omitted for clarity. Note that KLB binds closer to the PTC than PHZ. (**C, D**) Close-up view of PHZ (C) and KLB (D) interactions with the loops of proteins uL4 and uL22 forming the NPET constriction site. Superposition of the loops of proteins from *E. coli* and *T. thermophilus* is shown. Note that sterical clashes occur only with the side chains in the loops of proteins uL4/uL22 from *T. thermophilus*. (**E**) Multiple sequence alignment of the homologous parts of the ribosomal protein uL4 from several bacterial species. Red boxes highlight the amino acid residues of uL4, which either clash with the PHZ (His69 in *Tth*) or interact with it in our structure (Arg61 in *Eco*). (**F**) Comparison of the growth on the agar plates without PHZ (left) and with 10 μM PHZ (right) for *S. meliloti* Sm1021 and its derivatives expressing plasmid copies of the WT or the mutant *rplD* gene, which encodes for the ribosomal protein uL4. Note that overexpression of *Tth*-like version of uL4 protein in *S. meliloti* confers resistance to PHZ.

## DISCUSSION

The newly discovered antibiotic phazolicin represents the second known example of a linear azole-containing peptide with a known mechanism of action and mode of binding to its intracellular target, the ribosome. Although the PHZ and the previously described LAP klebsazolicin both inhibit the elongation step of prokaryotic translation through the obstruction of the NPET of the large ribosomal subunit, the mode of binding of PHZ to the bacterial ribosome is significantly different from that of KLB. Unlike KLB, which does not appear to have extensive intramolecular interactions, four out of eight azole rings of PHZ form continuous π-π-stacking system (**Figure S5D**), which apparently improves the rigidity and stability of the antibiotic in its binding site. Moreover, PHZ contains several positively charged side chains, which provide additional anchoring on the ribosome through the electrostatic interactions with the negatively charged 23S rRNA (**Figure 3E**), and which were absent in KLB. Finally, PHZ appears to be the first ribosome-targeting antibiotic with a pronounced bacterial species-specific mode of action, which is directly related to its mode of interaction with the molecular target. PHZ can bind to the ribosomes and act upon those species whose ribosomal protein uL4 contains glycine (or non-bulky side chain amino acid residues) in the position equivalent to Gly64 in *E. coli* that is located at the tip of the extended loop of this protein. The residue equivalent to Gly64 of the *E. coli* protein uL4 is not conserved in other bacterial species. The majority of bacteria (including *E. coli*) possess glycine at this position (**Figure 5E**). However, others (e.g., *T. thermophilus*) contain histidine that renders them naturally resistant to this antibiotic due to a steric hindrance with PHZ molecule. There are many ribosome-targeting antibiotics that, as PHZ (see **Table 1**), work on one group of bacteria and do not work on the others, however, this phenomenon is usually not referred to as “species-specificity of action” because it is due to the ability of a drug to get inside the bacterial cell rather than a differential activity on its target.

To our knowledge, PHZ is the first example of a bacterial ribosome-targeting inhibitor whose activity depends not only on its ability to penetrate inside the cell but also on the fine structure of its target, the 70S ribosome, regardless of its methylation. Moreover, PHZ is the second known example of characterized RiPPs from the order *Rhizobiales*. Owing to the ability of nitrogen fixation and establishing symbiotic relationships with leguminous plants, rhizobia are considered as an agriculturally important group of bacteria that have been extensively studied for decades. However, even with dozens of full-genome sequences, the number of rhizobial natural products that were characterized to date remains extremely small in comparison to such fruitful groups as the *Actinobacteria* or the *Enterobacteriales*. At the same time, genome mining reveals significant potential for the discovery of novel RiPPs in this group of bacteria. Of particular interest are the genes associated with the biosynthesis of LAPs and Nif11/NHLP-containing peptides^33^ present in multiple rhizobial genomes.

Characterization of PHZ together with the identification of biosynthetic gene clusters of its predicted homologs in the genomes of other *Rhizobiales* expands the diversity of LAPs and illustrates how different RiPPs can share similar enzymatic machinery and, at the same time, can exhibit significantly different mechanisms of binding and action upon the target. The narrow-spectrum antibacterial activity of PHZ against rhizobia may serve as a starting point for the development of biocontrol agents, which could be used to combat for example plant-pathogenic *Agrobacterium* strains, or rhizobium inoculants for legume crops that can outcompete endogenous rhizobium strains.

## MATERIALS AND METHODS

### Bacterial strains and growth conditions

The following bacterial strains were used in the present study: *Rhizobium sp.* Pop5, *R. leguminosarum* bv. *viciae* 3841, *R. leguminosarum* bv. *phaseoli* 4292, *R. leguminosarum* bv. *phaseoli* RCR 3622, *R. etli* DSM 11541, *R. tibethicum* DSM 21102, *Agrobacterium tumefaciens* C58C1, *A. rhizogenes* ARQUA1, *Sinorhizobium meliloti* Sm1021, *S. fredii* HH103*, S. medicae* WSM419, *Mesorhizobium ciceri*, *M. loti* MAFF303099, *M. thianshanense* HAMBI 3372, *Azorhizobium caulinodans* ORS571, *Erwinia amylovora, Pantoea ananatis* PA4*, Pseudomonas putida* KT2440*, Pseudomonas fluorescens* Pf-5*, Bacillus subtilis* 168*, Bacillus cereus* ATCC 4342*, Arthrobacter* sp. ATCC21022, *Microtetraspora glauca* NRRL B-3735, *Escherichia coli* DH5α*, E. coli* HB101, *E. coli* MRE600, *E. coli* BW25113, and *E. coli* BW25113Δ*tolC*. We used RM medium (per 1L: 10 g mannitol, 0.5 g K2HPO4, 0.2 g MgSO4, 0.1 g NaCl, 1 g yeast extract pH 6.8) for cultivation of various species of the *Rhizobium* genus (**Table 1**). Other rhizobia (including species of the genera *Mesorhizobium, Sinorhizobium, Agrobacterium* and *Azorhizobium*) were cultivated in YEB medium (per 1 L: 5 g beef extract, 5 g peptone, 5 g sucrose, 1 g yeast extract, 0.2 g MgSO4, pH 7.5) (**Table 1**). All rhizobia strains were cultivated at 28°C. Growth of all other strains was performed in LB medium (per 1 L: 5 g NaCl, 10 g tryptone, and 5 g yeast extract) at 28°C or 37°C (for *E. coli* strains). When necessary, antibiotics were used at concentrations of 100 μg/ml for kanamycin (for *S. meliloti* Sm1021), 50 μg/ml for kanamycin (for *E. coli* strains), 25 μg/ml for chloramphenicol, and 500 μg/ml for streptomycin.

### Molecular cloning procedures

Primers used for DNA amplification are listed in **Table S1**. The *rplD* gene was amplified from *Sinorhizobium meliloti* Sm1021 genomic DNA using rplD_pSRK_GA_F and rplD_pSRK_GA_R primers and cloned into the pSRK (Kan) broad host range vector^34^ following the Gibson assembly protocol (NEB). Two mutant variants of the *rplD* gene were obtained through PCR-based site-specific mutagenesis with primer pairs rplD_G68H_F/rplD_mut_R and rplD_G68H+K65A_F/rplD_G68H+K65A_R and cloned into the pSRK plasmid as well. The *phzE* gene was amplified from *Rhizobium sp.* Pop5 genomic DNA using phzE_pSRK_GA_F and phzE_pSRK_GA_R primers and cloned into pSRK. The transformation of *Sinorhizobium meliloti* Sm1021 with pRSK and pSRK-derived vectors was performed by triparental mating using pRK600 as a helper plasmid.

### PHZ production and purification

*Rhizobium sp.* Pop5 was cultivated for 24 hours in liquid RM medium (750 ml) with shaking (180 rpm) at 28°C. Next, the cells were pelleted (14,000 rpm, 30 min, 4°C) and cultivation medium supplied with 0.1% trifluoroacetic acid (TFA) was loaded onto Agilent HF Bond Elut LRC-C18 Cartridge (1 g) pre-equilibrated with 0.1% TFA. The cartridge was then extensively washed with water followed by 50 ml of 10% acetonitrile (MeCN). The peptide fraction was eluted using 10 ml of 30% MeCN, dried out with centrifugal evaporator (GeneVac EZ-2), and dissolved in 500 μL of 100% dimethyl sulfoxide (DMSO). The obtained extract was then subjected to reverse phase HPLC purification on a Luna PREP C18 column (21.2×250 mm, 10μm particle size). The column was pre-equilibrated using 0.1% TFA (Buffer A). The following gradient of 100% MeCN (Buffer B) was used for the separation and elution of the peptides: 0-5 min 0% B, 5-12 min 0-19% B, 12-22 min 19-22% B, 22-27 min 22-70% B, 27-33 70% B. The detection was performed at 254 nm, which is a characteristic wavelength for azole-containing compounds. The collected fractions were then checked with MALDI-ToF MS and those containing the compounds of interest were dried under vacuum, dissolved in 100% DMSO and stored at −20°C until further use.

### High-resolution MS and ESI-MS-MS analysis

One microgram of peptides in a volume of 1-4 μL was loaded onto the Acclaim μ-Precolumn (0.5 mm х 3 mm, 5 μm particle size, Thermo Scientific) at a flow rate of 10 μL/min for 4 min in an isocratic mode of Mobile Phase C (2% MeCN, 0.1% formic acid). Then the peptides were separated with high-performance liquid chromatography (HPLC, Ultimate 3000 Nano LC System, Thermo Scientific, Rockwell, IL, USA) in a 15-cm long C18 column (Acclaim^®^ PepMap™ RSLC inner diameter of 75 μm, Thermo Fisher Scientific, Rockwell, IL, USA). The peptides were eluted with a gradient of buffer B (80% MeCN, 0.1% formic acid) at a flow rate of 0.3 μL/min. The total run time was 60 minutes, which included an initial 4 min of column equilibration with buffer A (0.1% formic acid), then gradient from 5 to 35% of buffer B over 35 min, then 6 min to reach 99% of buffer B, flushing 10 min with 99% of buffer B and 5 min re-equilibration to buffer A.

MS analysis was performed at least in triplicate with a Q Exactive HF mass spectrometer (Q Exactive HF Hybrid Quadrupole-OrbitrapTM Mass spectrometer, Thermo Fisher Scientific, Rockwell, IL, USA). The temperature of the capillary was 240°C and the voltage at the emitter was 2.1 kV. Mass spectra were acquired at a resolution of 120,000 (MS) in a range of 300−1500 *m/z*. Tandem mass spectra of fragments were acquired at a resolution of 15,000 (MS/MS) in the range from 100 *m/z* to 1200 m/z value determined by a charge state of the precursor, but no more than 2000 *m/z*. The maximum integration time was 50 ms and 110 ms for precursor and fragment ions, respectively. Automatic gain control (AGC) target for precursor and fragment ions were set to 1*10^6^ and 2*10^5^, respectively. An isolation intensity threshold of 50,000 counts was determined for precursor’s selection and up to top 20 precursors were chosen for fragmentation with high-energy collisional dissociation (HCD) at 29 NCE (Normalized collision energy). Precursors with a charge state of +1 and more than +5 were rejected and all measured precursors were dynamically excluded from triggering of a subsequent MS/MS for 20 s.

### MALDI-ToF-MS and MS-MS analysis

MALDI-MS mass spectra were recorded in a positive ion measurement mode using Ultraflextreme MALDI-ToF-ToF-MS instrument (Bruker Daltonics, Germany). The spectra were obtained in reflecto-mode with the accuracy of measuring the monoisotopic m/z ratio up to 0.1 Da. The fragmentation spectra were obtained using the Lift mode, for which the accuracy of measuring the daughter ions was within 0.2 Da. Sample aliquots were mixed on a steel target with a 30 mg/ml 2,5-dihydroxybenzoic acid in 0.5% trifluoroacetic acid and a 30% MeCN-water solution (Sigma-Aldrich).

### Determination of minimal inhibitory concentrations (MICs)

Determination of MICs was performed in 96-well plates in different media for different bacterial strains (**Table 1**) using a microdilution assay. In each row, the next well contained two times less antibiotic than the previous one. Two different concentration ranges were tested: 128 μM to 0.5 μM for the strains less susceptible to PHZ; and 8 μM to 31.25 nM for the strains more susceptible to PHZ; a well without antibiotic was used as a control. Initially each well contained 100 μL of media. Actively growing bacteria were added to the wells up to the concentration of approximately 5*10^6^ cells/mL. The plates were incubated for 48 hours at 28°C with shaking (180 rpm). MIC was defined as the concentration of PHZ in the well, at which no bacterial growth was observed, while the two-fold lower concentration of PHZ was not abolishing growth of bacteria.

### Testing of PHZ activity in vivo

For the *in vivo* bioactivity test (**Figure 2A**), the reporter strain BW25113-pDualrep2 and BW25113*ΔtolC*-pDualrep2 was used as previously described^19^. Briefly, 2 μL of solutions of tested compounds were applied onto the agar plate that already contained a lawn of the reporter strain. 10 mM PHZ, KLB, erythromycin, and 20 μM levofloxacin solutions were used. After overnight incubation of the plate at 37°C, it was scanned by ChemiDoc (Bio-Rad) using “Cy3-blot” settings for RFP and “Cy5-blot” for the Katushka2S protein.

### In vitro translation inhibition assays

The inhibition of firefly luciferase synthesis by PHZ (**Figure 2B**) was assessed essentially as described previously^23^. Briefly, the *in vitro* transcribed firefly luciferase mRNA was translated in the *E. coli* S30 Extract System for Linear Templates (Promega). Reactions containing 200 ng of mRNA and 0.1 mM of D-luciferin were carried out in 5 μL aliquots at 37°C for 15 minutes and the activity of *in vitro* synthesized luciferase was measured by VictorX5 (PerkinElmer) every 30 seconds.

To test the ability of PHZ to act upon the *E. coli* vs. *T. thermophilus* ribosomes (**Figure S3**), we used *in vitro* PURExpress translation system (New England Biolabs) reconstituted from purified components and supplied with either the WT 70S ribosomes from *E. coli* or the reconstituted hybrid 70S ribosomes containing purified 30S subunits from *E. coli* and purified 50S subunits from *T. thermophilus*. Translation of superfolder green fluorescent protein (sfGFP) was carried out according to the manufacturer’s protocol. The assembled reactions (10 μL) were supplemented with 100 ng of sfGFP mRNA. PHZ, KLB, or CHL (chloramphenicol) antibiotics we added to a final concentration of 50 μM when needed. The reactions were placed in a 384-well black-wall plate and the progression of the reactions was monitored over 3 hours by a TECAN microplate scanner.

### Toe-printing analysis

Toe-printing analysis was carried out essentially as previously described^35^, using *trpL*-2Ala mRNA as a template for protein translation. *trpL*-2Ala template was prepared by PCR from pDualrep2 plasmid with following oligonucleotides: F TGTAATACGACTCACTATAGGGAATTTTTCTGTATAATAGCCGCGGAAGTTCACGTAAAAAGGGTATCGACAATGAA and R GCGTTAAGGCTATGTACGGGTATCTGATTGCTTTACG, GCGTTAAGGCTATGTAC was used for reverse transcription. The concentrations of the tested compounds are shown above each lane in **Figure 2E**.

### Cryo-electron microscopy (cryo-EM) structure determination

*Ribosome preparation: E. coli* strain MRE600 overnight cultures were diluted 1:100 into 3L of LB. After growth to mid-log phase (OD600=0.6), cells were cooled on ice for 3 min, pelleted, and washed with buffer A (20 mM Tris-HCl pH 7.5, 100 mM NH4Cl, 10 mM MgCl2, 0.5 mM EDTA, 2mM DTT). Cells were resuspended in 100 mL buffer AS (buffer A with 0.15 M sucrose) and lysed with two passes through an Emulsiflex-C3 homogenizer at 18000 psi. Lysate was clarified by centrifugation for 30 min at 18000 rpm in a JA-20 rotor (Beckman-Coulter). The supernatant was layered over a sucrose cushion of 24mL buffer B (buffer A with 500mM NH4Cl) with 0.5M sucrose and 17mL buffer C (20 mM Tris-HCl pH 7.5, 60 mM NH4Cl, 6 mM MgCl2, 0.5 mM EDTA, 2 mM DTT) with 0.7M sucrose in ti-45 tubes (Beckman-Coulter). Ribosomes were pelleted in a Ti-45 rotor at 27000 rpm for 16 hours at 4°C. Crude ribosome pellets were resuspended in disassociation buffer (buffer C with 1 mM MgCl2) and clarified via centrifugation at 15000g for 10 min. The cleared supernatant was layered over 15-40% sucrose gradients in disassociation buffer and spun 28000 rpm for 16 hours at 4°C in a SW-32 rotor (Beckman-Coulter). Gradients were fractionated with an ISCO fractionation system and 30S + 50S peaks were combined. The combined subunits were concentrated in 15 mL Millipore 100k molecular weight cutoff spin filter and buffer exchanged with reassociation buffer (buffer C with 10 mM MgCl2). Ribosomes were incubated 45min at 37°C, layered onto 15-40% sucrose gradients in reassociation buffer, and spun 27000 rpm for 15 hours at 4°C in an SW-32 rotor. Gradients were fractionated as before, 70S peaks were collected, and ribosomes were concentrated and washed with buffer C in an Amicon stirred cell filtration system using 100k cutoff filters. Ribosomes were stored in aliquots at −80°C.

#### Cryo-EM sample preparation

Roughly 4-6 μM PHZ was incubated with 100 nM ribosomes in RC buffer (50 mM Hepes pH 7.5, 150 mM KOAc, 12 mM MgOAc, 7 mM β-mercaptoethanol) for 30 min. at 37°C. Complexes were deposited in 4-μL aliquots on 300 mesh Quantifoil UltraAuFoil R1.2/1.3 grids that were topped with a layer of amorphous continuous carbon and glow-discharged for 15 seconds with a Pelco SC-6 sputter coater. After incubation period of about a minute, the excess sample was washed off on drops of RC-LS buffer (RC with 25mM KOAc). Samples were blotted and plunge-frozen in liquid ethane using an FEI Vitrobot with blot force 6, humidity 100%, and 20°C or 4°C depending on the freezing session.

#### Data collection

Images were collected in two separate sessions on an FEI Titan Krios Microscope operated at 300 keV with a GIF energy filter and GATAN Summit K2 direct electron detector in super-resolution mode. Images were collected at 215,000x magnification for a pixel size of 0.56 Å (0.28 Å super-resolution). Automated movie collection was performed with SerialEM^36^ over the defocus range −0.6 to −2.0 μm, and Focus software^37^ was used to monitor data collection. The total dose was 29.97 e^−^/Å^2^ for session 1 and 29.83 e^−^/Å^2^ for session 2, each over 30 frames.

#### Cryo-EM data analysis

Cryo-EM data processing was done using RELION 3.0 software^38^ (**Figure S9; Table S2**). The two datasets were processed separately until the final round of map refinement, for which the best particles from both sets were combined. Movies were motion-corrected and dose-weighted using RELION’s motion correction algorithm. Contrast transfer function (CTF) parameters were estimated using CTFFind4^39^ and micrographs with poor CTF fit, as determined by visual inspection, were sorted out. Particles were picked automatically with the Laplacian-of-Gaussian method and subjected to several rounds of careful 2D class-based cleaning before an initial round of 3D auto-refine. CTF refinement with per-particle defocus and beam tilt correction was performed, and particles were further sorted through 3D classification without alignment. Another round of 3D auto-refine was performed on the best 3D classes with focused refinement on the 50S subunit. This was followed by one round of Bayesian polishing^38^ and another round of CTF refinement, which were each followed by another round of 3D auto-refine with a mask around the 50S subunit. Particles from the two sessions were then pooled for the final round of 3D auto-refine. A charge density map was calculated in Chimera^40^ as previously described^41^. Post-processing was also performed in RELION to generate a Fourier shell correlation (FSC) curve. All refinement procedures were run on 2x binned particles because of the small pixel size.

*Model building.* Assembly #2 from the PDB entry 4YBB^42^ was used as a starting ribosome model. Only the 50S subunit was subjected to real-space refinement in PHENIX^43^, because charge density corresponding to the 30S subunit was relatively poor after focused refinement on the 50S, where PHZ binds. Ions and water molecules were removed from the starting model prior to real-space refinement. The atomic model for the PHZ was built as follows: first, baton building in COOT^44^ was used to trace the main chain; next, the resulting polyalanine chain was mutated to the unmodified PHZ sequence after visual search for the characteristic features in the map; next, oxazole and thiazole modifications were built in Avogadro^45^ version 1.2.0 (http://avogadro.cc) based on the density. Finally, CIF restraints for the oxazole and thiazole moieties were created based on the structural data from^46^.

All figures showing atomic models were generated using the PyMol software (www.pymol.org). USCF Chimera was used for EM density maps shown in Figs. S5 and S9.

## ACCESSION NUMBERS

Coordinates and charge density map were deposited in the RCSB Protein Data Bank with accession code XXX for the *E. coli* 50S ribosomal subunit in complex with PHZ.

## AUTHOR CONTRIBUTIONS

All authors interpreted the results. D.Y.T, Y.S.P., J.D.H.C., and K.S. wrote the manuscript.

## ACKNOWLEDGMENTS

We thank Dr. Viktor Zgoda for the invaluable help with high-resolution spectra and ESI-MS-MS spectra acquisition and Dr. Esperanza Martinez-Romero who provided the strain *Rhizobium* sp. Pop5. We thank all members of the K.S., J.H.D.C., and Y.S.P. laboratories for important discussions and critical feedback. We also want to thank Dr. Dmitry Ghilarov and Dr. Svetlana Dubiley for critical reading of the manuscript and valuable suggestions.

This work was supported by RSF grant number 19-14-00266 to Svetlana Dubiley, RSF grant number 18-44-04005 to I.A.O. (used for the toe-printing assays and *in vivo* (pDualrep2 reporter) and *in vitro* activity tests), by Illinois State startup funds [to Y.S.P.], National Institutes of Health [R01-GM132302 and R21-AI137584 to Y.S.P., R01-GM114454 to J.H.D.C and F.R.W] and by the National Science Foundation Graduate Research Fellowship Program under Grant number 1106400 [to Z.L.W.]. Access to MALDI-MS facilities was through the framework of the Lomonosov Moscow State University Development Program PNR 5.13.

