## Supplementary material for "Phazolicin – a Novel Thiazole/Oxazole-Modified Peptide Inhibiting the Bacterial Ribosome in a Species-Specific Way": Supplimentary Information

**This file includes:**

I. Supplementary Figures 1 to 9 with legends;
II. Supplementary Movie 1 legend;
III. Supplementary Tables 1 and 2;
IV. Supplementary References.

### I. SUPPLEMENTARY FIGURES

A

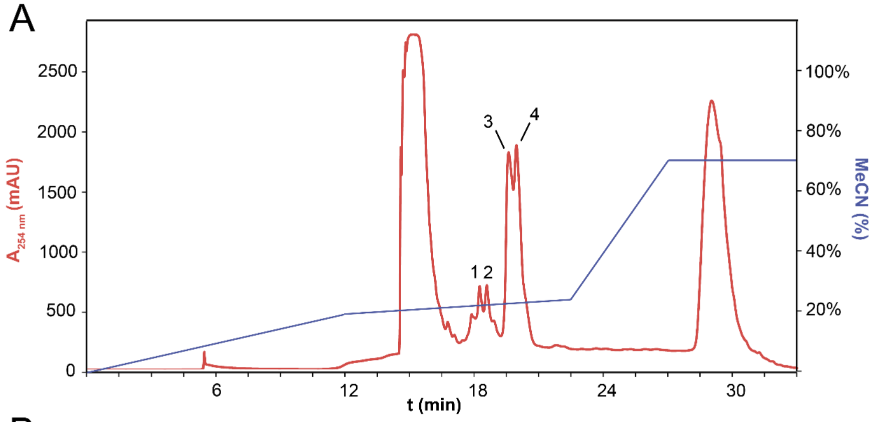


B

| | HPLC peak | Formula | Mass (av.) | Number of cycles | Structural features |
|---|---|---|---|---|---|
| Major forms | 3 | ATCARCDSSSRCGASGKSSGSASSIST | 2364.5 | 8 | Phasolicin (PHS) |
| | 4 | AATCARCDSSSRCGASGKSSGSASSIST | 2435.5 | 8 | Alternative leader peptide cleavage site |
| | 4 | TAATCARCDSSSRCGASGKSSGSASSIST | 2536.6 | 8 | |
| Minor forms | 1 | ATCARCDSSSRCGASGKSSGSASSIST | 2384.5 | 7 | Smaller number of cycles in the product |
| | 1 | ATCARCDSSSRCGASGKSSGSASSIST | 2404.5 | 6 | |
| | 1 | ATCARCDSSSRCGASGKSSGSASSIST | 2424.5 | 5 | |
| | 2 | AATCARCDSSSRCGASGKSSGSASSIST | 2455.5 | 7 | Alternative leader peptide cleavage site + smaller number of cycles |
| | 2 | TAATCARCDSSSRCGASGKSSGSASSIST | 2556.6 | 7 | |
| | 1,2 | ATCARCDSSSRCGASGKSSGSASS | 2083.4 | 7 | C-terminal degradation of the peptide + alternative cleavage |
| | 1,2 | AATCARCDSSSRCGASGKSSGSASS | 2154.4 | 7 | |

C

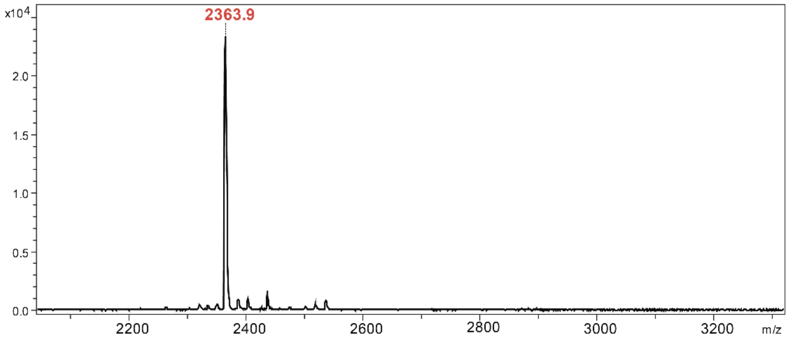


**Supplementary Figure 1 | Isolation and purification of PHZ.** (**A**) HPLC profile of the crude extract from RM medium after *Rhizobium sp.* Pop5 cultivation. Shown are the 254-nm absorbance curve (red trace) together with the MeCN-gradient profile (blue trace). Note that azole-containing peptides absorb UV at 254-nm wavelength. Each of the HPLC peaks was assigned a number and subsequently analyzed using MS. (**B**) Characteristics of PHZ-related compounds from the medium extracts. Major forms (HPLC peaks 3 and 4) are indicated with the dark blue background. Cyclized residues are highlighted in red in the peptide sequences. (**C**) MS spectra of HPLC peak 3. Molecular mass of 2363.9 $[M+H]^+$ corresponds to PHZ.

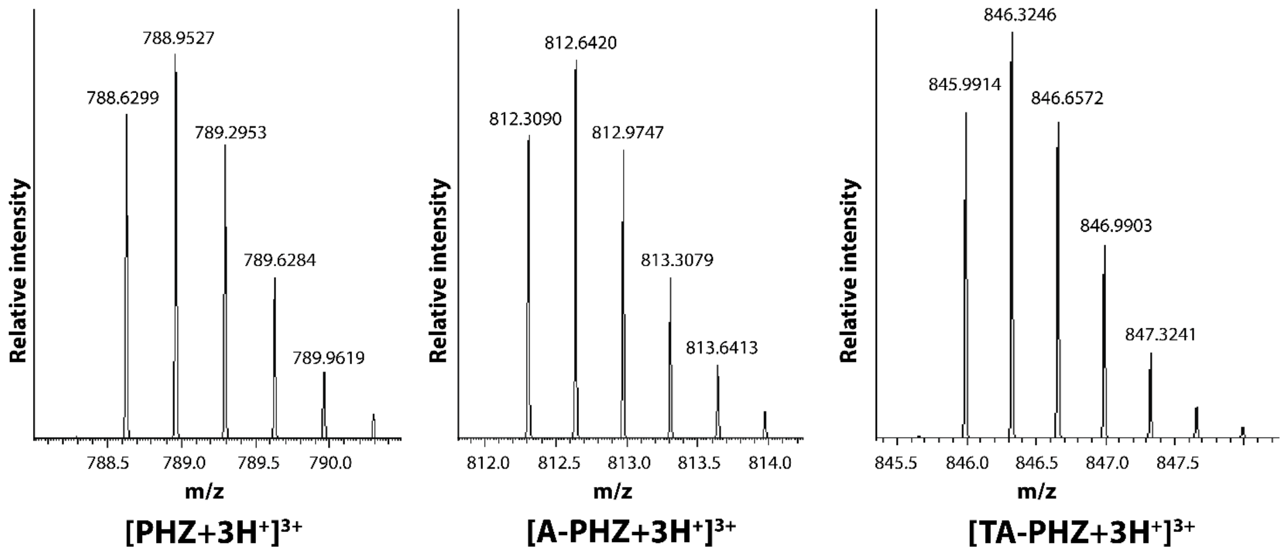


| | | | |
|---|---|---|---|
| **Calculated MW:** | 2362.8651 | 2433.9022 | 2534.9499 |
| **Experimental MW:** | 2362.8679 | 2433.9052 | 2362.9524 |
| **Mass error (ppm):** | 1.18 | 1.23 | 0.98 |

**Supplementary Figure 2 |** High-resolution ESI-MS spectra of the three major compounds related to PhzA precursor peptide identified in the *Rhizobium sp.* Pop5 cultivation medium (mature PHZ and two forms with additional amino acid residues at the N-terminus, resulting from alternative leader cleavage). The m/z values for each peak are indicated. The calculated mass, experimentally obtained mass, and mass error in ppm are shown for each compound.

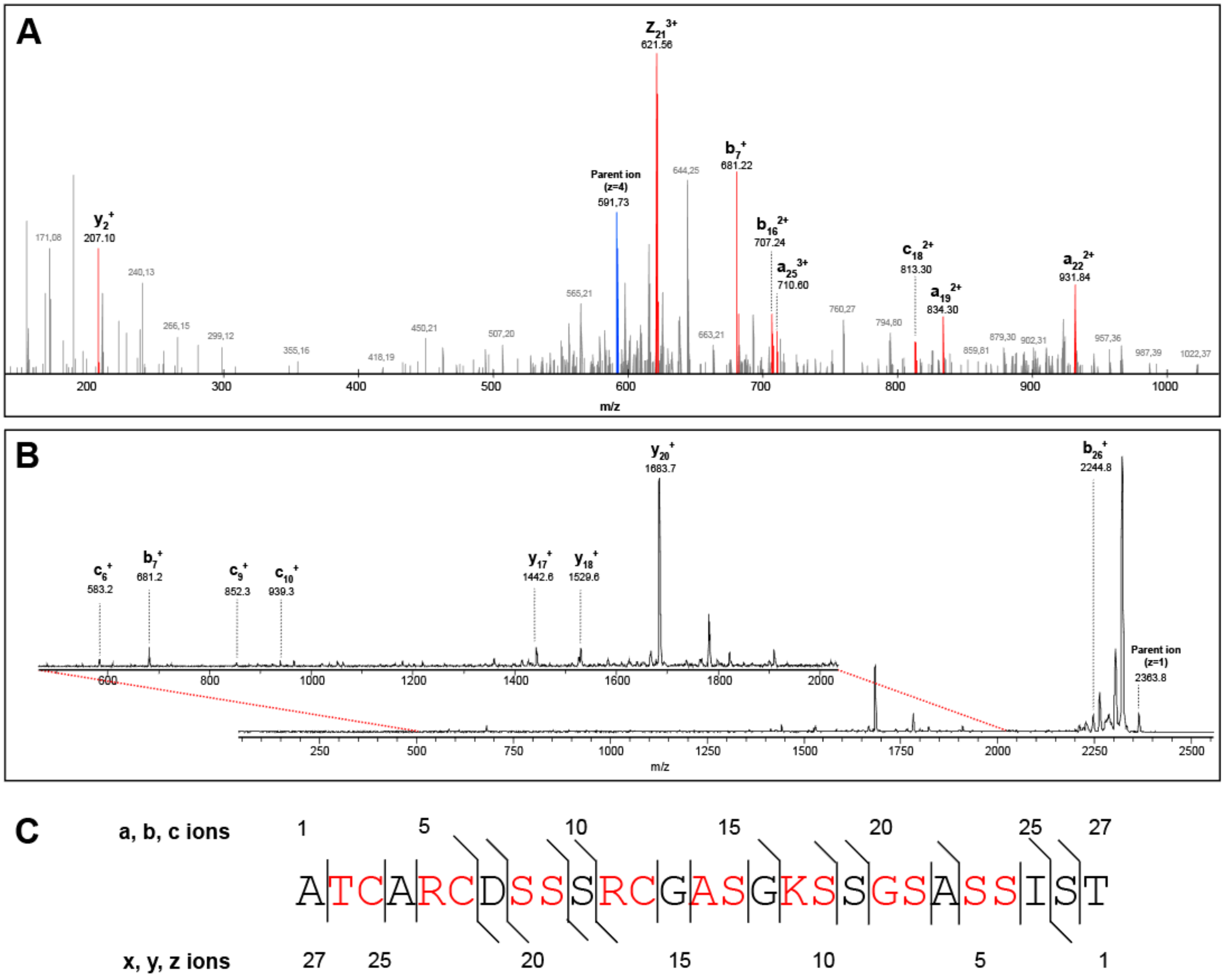


**Supplementary Figure 3 | MS/MS analysis of PHZ. (A)** ESI-MS/MS spectrum for the ion 591.73 $[M+4H^+]^{4+}$. The peaks corresponding to single fragmentation events in the structure of the compound are shown in red and labeled. **(B)** MALDI-MS/MS spectrum for the ion 2363.8 $[M+H^+]^+$. The range of m/z between 500 and 2050 is shown in detail separately. **(C)** Amino acid sequence of PHZ. The fragments detected in MS/MS analysis are indicated with slashes, residues proposed to be involved in azole cycle formation are red, numeration of ions is shown above (a,b,c series) and below (x, y, z series) the sequence.

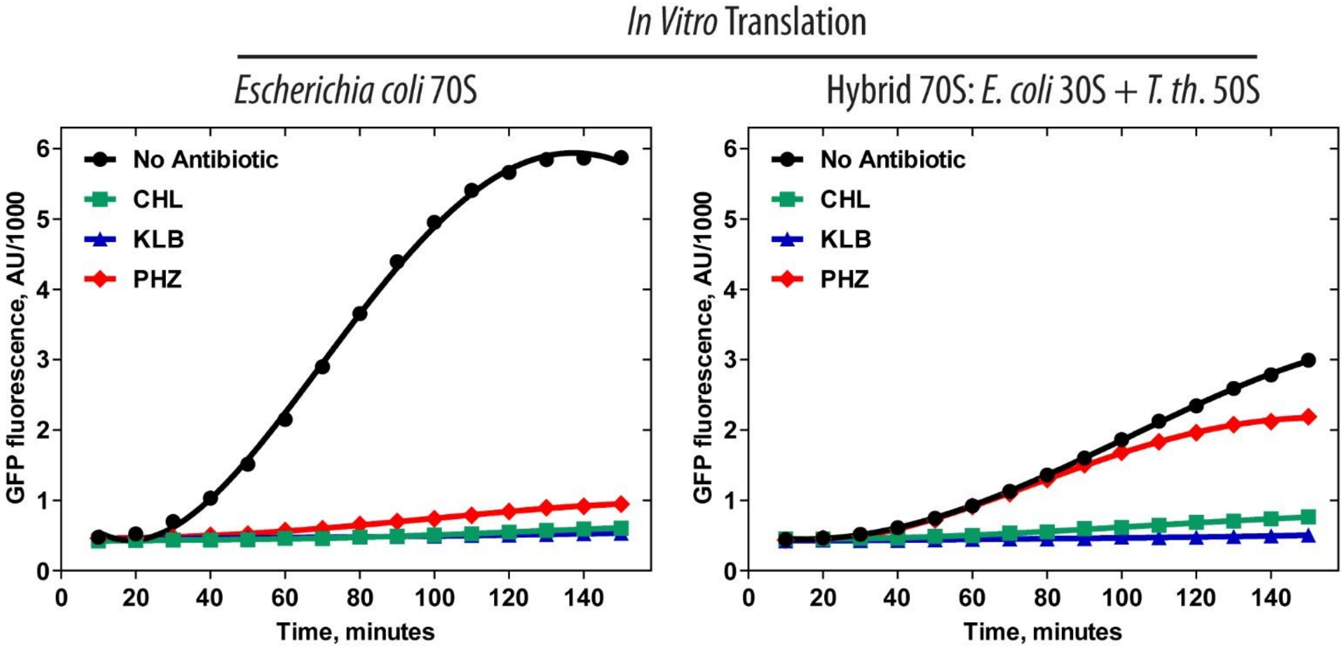


**Supplementary Figure 4 |** Effects of phazolicin (PHZ), chloramphenicol (CHL), and klebsazolicin (KLB) on *in vitro* synthesis of superfolder green fluorescent protein (sfGFP) using PURExpress system supplied with either the *E. coli* 70S WT ribosomes or hybrid ribosomes (*E. coli* 30S + *Th. thermophilus* 50S). CHL and KLB were used as positive controls. Final concentrations for all antibiotics were 50 µM. Note, that the overall rate of protein synthesis by the hybrid ribosome is ~2-fold lower than that of the WT 70S (black curves).

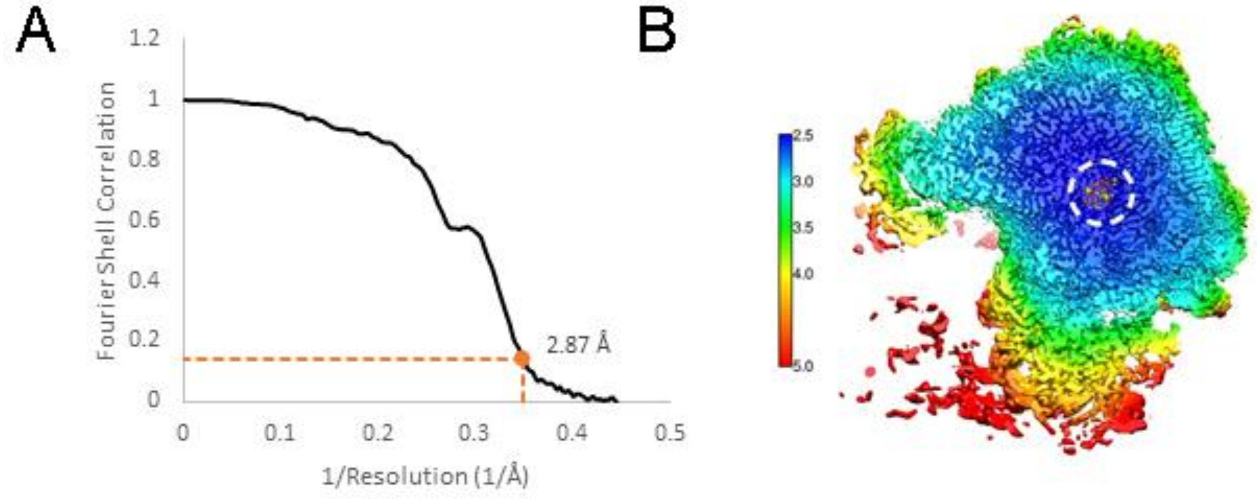


**Supplementary Figure 5 | Fourier shell correlation (FSC) curve for the 70S-PHZ cryo-EM structure.** (**A**) Based on the "gold-standard FSC" cutoff value 0.143, the overall resolution for the large ribosomal subunit in the 70S ribosome is 2.87 Å. (**B**) Slice through the 50S subunit showing local resolution limits. Note that the region of PHZ binding (indicated with the dashed white circle) is characterized by the highest local resolution. Color scale bar on the left is in Ångstroms. 30S ribosomal subunit is not shown and was not included in the final model because after the focused refinement on the 50S subunit the local resolution and connectivity of the map for the 30S subunit were poor.

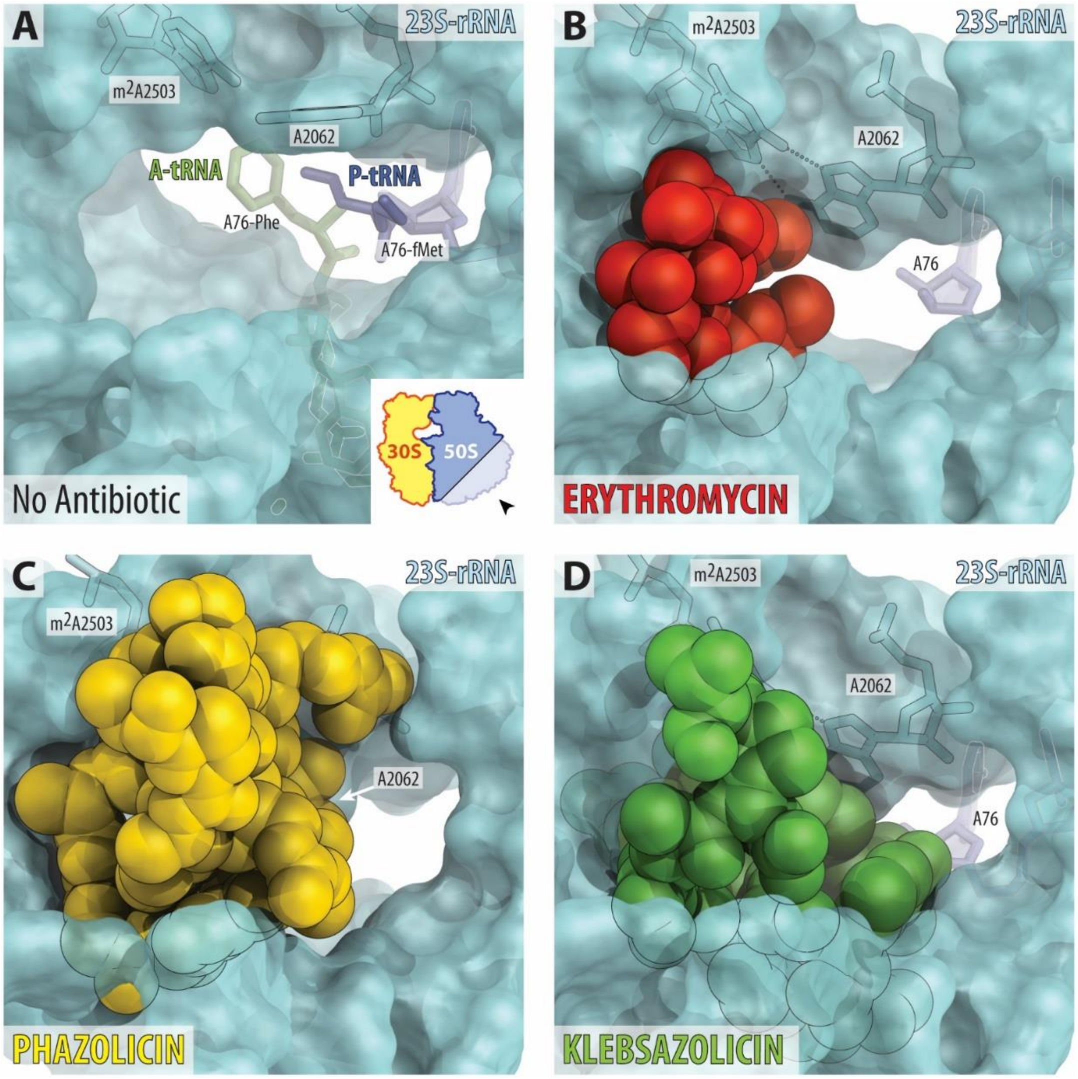


**Supplementary Figure 6 | Occlusion of the nascent peptide exit tunnel by antibiotics.** (**A**) The lumen of the nascent peptide exit tunnel of the drug-free 70S ribosome (PDB entry 4Y4P[1]). The view is from the wide-open part of the tunnel onto the PTC as indicated by the inset. Nucleotide A76 of the P-site tRNA is shown in dark blue. A-site tRNA is not visible in this view, however, the location of its aminoacylated moiety is shown. Note that nucleotide A2062 of the 23S rRNA is pointed toward the viewer and is not involved in Hoogsteen-edge base pairing with the nucleotide m$^2$A2503 in the wall of NPET. (**B**, **C**, **D**) Occlusion of the nascent peptide exit tunnel by ERY (B), PHZ (C), and KLB (D). Structures of ERY and KLB are from PDB entries 6ND6[2] and 5W4K[3], respectively. Note that, unlike ERY and similar to KLB, PHZ almost completely occludes the lumen of the peptide exit tunnel. Also note that binding of ERY, PHZ, or KLB causes characteristic rotation of nucleotide A2062 by more than ninety degrees away from the viewer to form Hoogsteen base-pair with the m$^2$A2503 of the 23S rRNA.

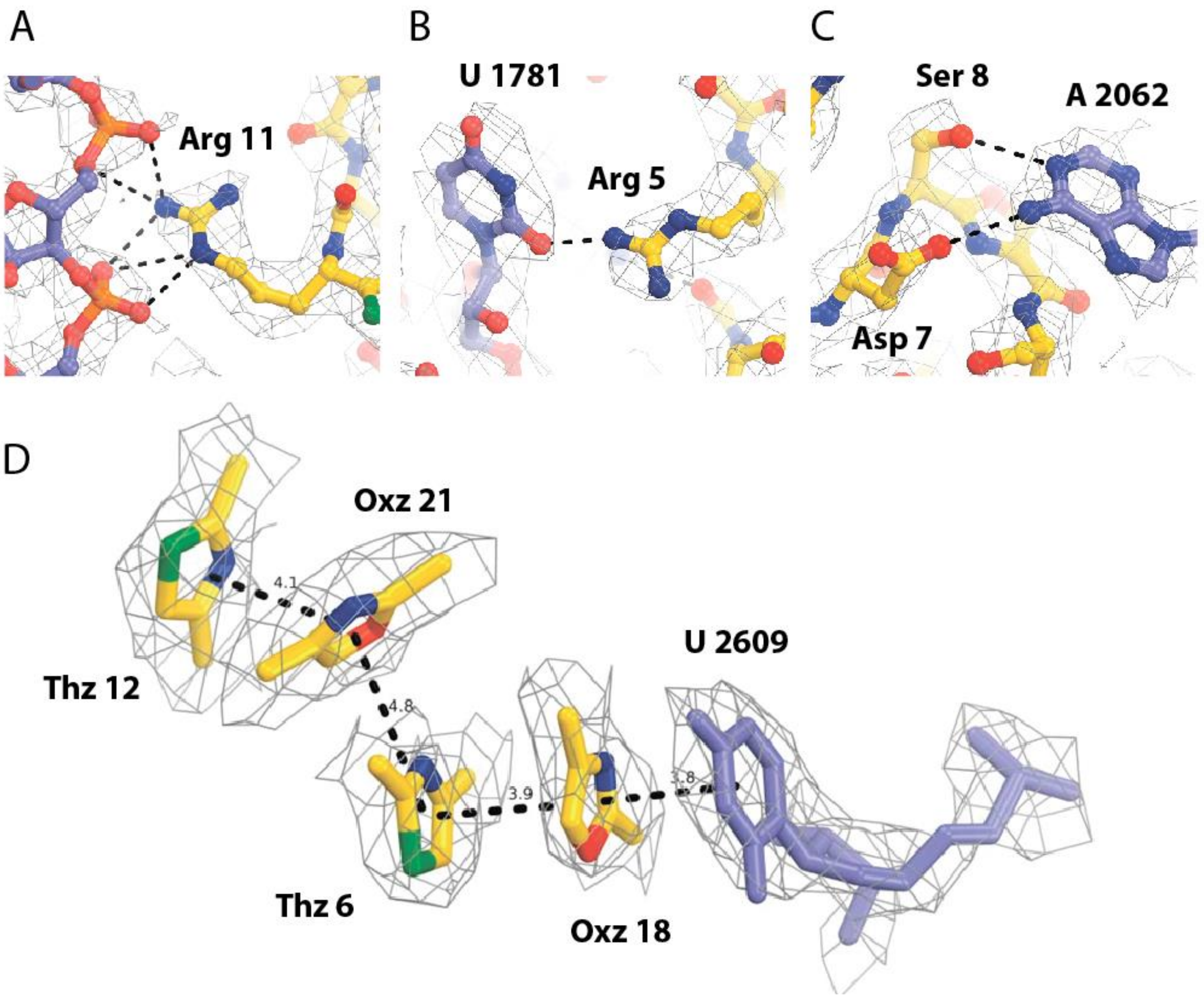


**Supplementary Figure 7 | PHZ contacts with the ribosome.** (**A-C**) Close-up views of the individual PHZ contacts with the nucleotides of the 23S rRNA. Hydrogen bonds are shown as dashed lines. (**D**) Close-up view of the intramolecular π-π-stacking system and its interaction with the nucleobase U2609 of the 23S rRNA. The directions of face-to-face and edge-to-face π-π stacking interactions are shown as dashed lines. Distances between the planes of aromatic heterocycles are shown in ångstroms (Å).

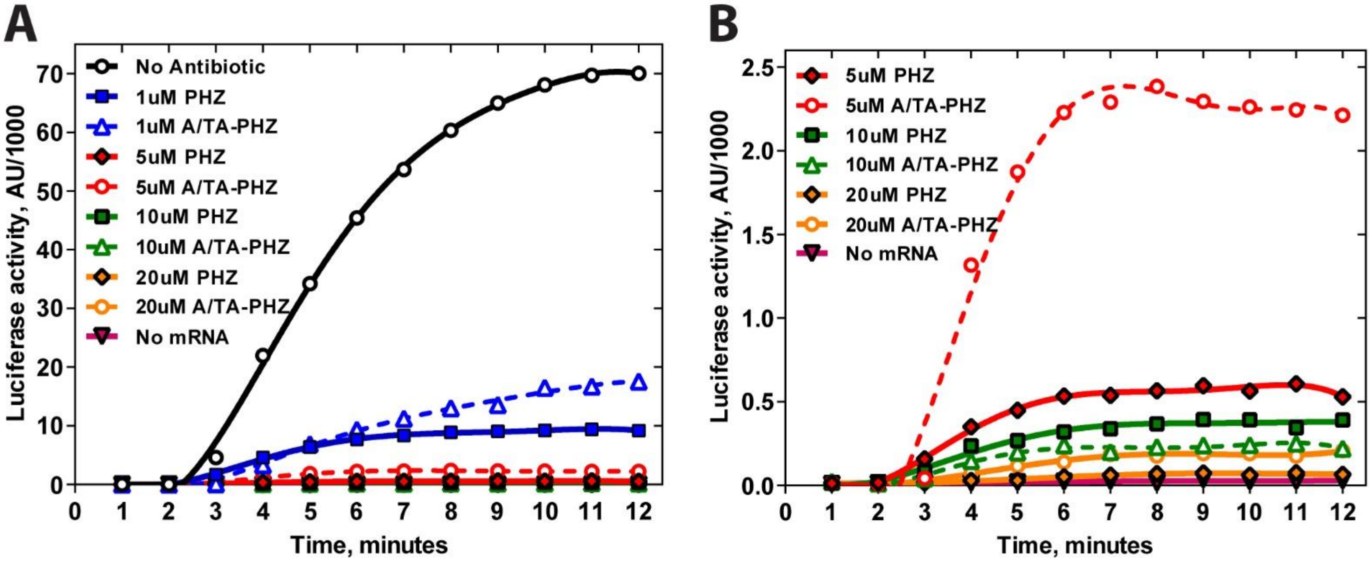


**Supplementary Figure 8 | Comparison of the *in vitro* translation inhibition activity of PHZ and its naturally occurring longer forms with additional amino acids at the N-terminus (A-PHZ and TA-PHZ).** Graphs corresponding to PHZ are shown as solid lines. Dashed lines correspond to the mixture of long forms of PHZ (A/TA-PHZ, HPLC peak 4, see Fig. S1A). **(B)** is a close-up view of **(A)** in the upper range of antibiotic concentrations (5-20 μM).

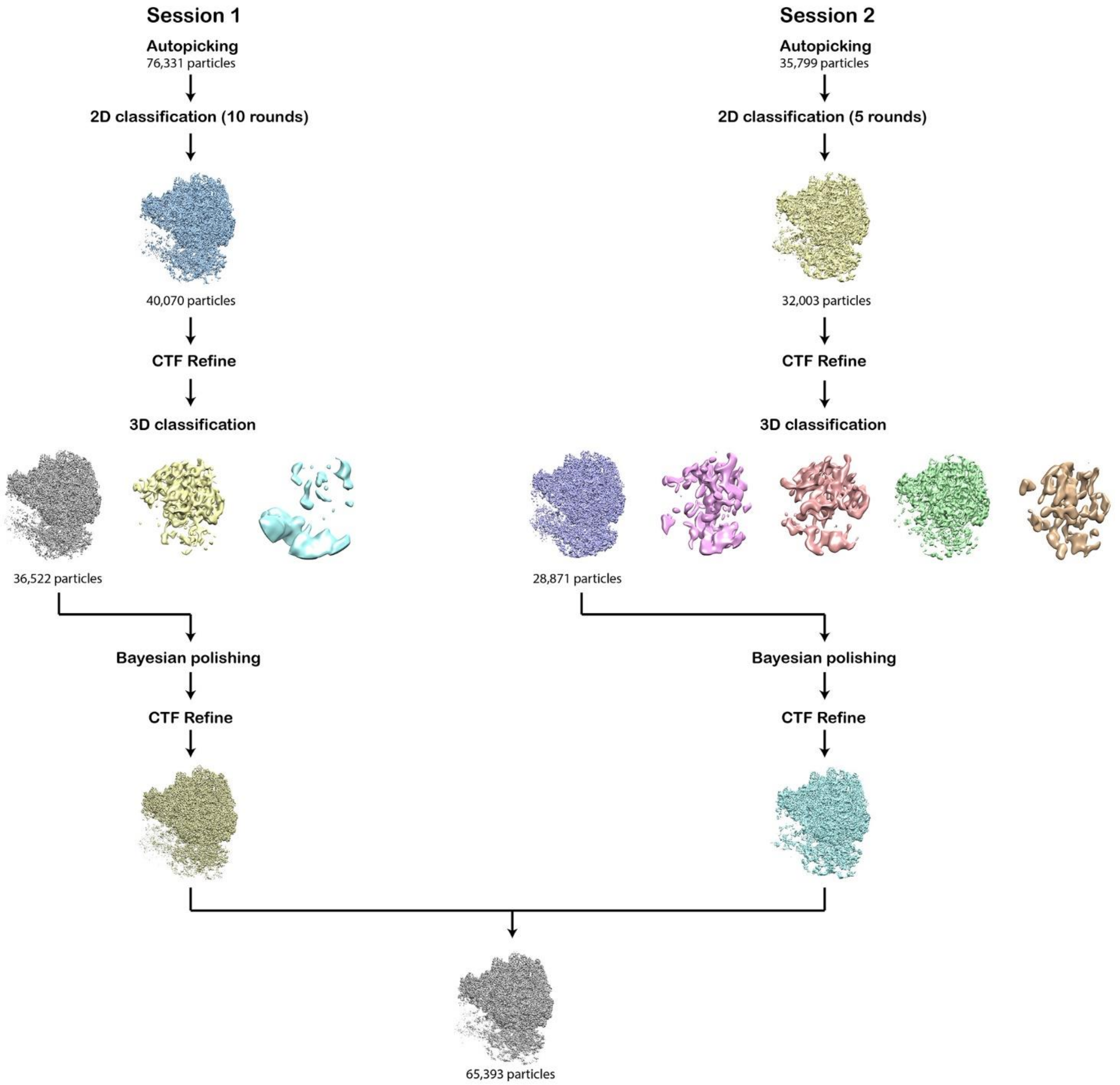


**Supplementary Figure 9 | Flowchart showing the steps of cryo-EM data processing in RELION.**

### II. SUPPLEMENTARY MOVIES

**Supplementary Movie 1 | PHZ functional site in the bacterial ribosome.** The movie shows: (1) zoom-out and (2) close-up views of the PHZ binding site in the large subunit of the *E. coli* ribosome; (3) details of PHZ interactions with the 23S rRNA in the PTC of the ribosome; (4) occlusion of the nascent peptide exit tunnel by the PHZ molecule.

### III. SUPPLEMENTARY TABLES

**Supplementary Table 1 | Nucleotide sequences of primers used for cloning.**

| Primer name | Primer sequence (5'-3') | Purpose |
|---|---|---|
| rplD_pSRK_GA_F | ATAACAATTTCACACAGGAAACAGCATATGGATCTCACCGTCAAAACCC | Molecular cloning of *rplD* gene into the pSRK plasmid (Gibson Assembly) |
| rplD_pSRK_GA_R | CGAGGTCGACGGTATCGATATCATTTGAACCGCTCCTCCAGAG | |
| pSRK_F_GA | ATGCTGTTTCCTGTGTGAAATTG | Amplification of the pSRK plasmid for Gibson Assembly cloning |
| pSRK_R_GA | TATCGATACCGTCGACCTCG | |
| rplD_G68H_F | TACAAGCAGAAGCATACGGGCCGCG | Site mutagenesis in *rplD* gene (Gly49His) |
| rplD_G68H_R | CGCGGCCCGTATGCTTCTGCTTGTACATC | |
| rplD_G68H+K65A_F | AGATGTACGCGCAGAAGCATACGGGCCGCG | Site mutagenesis in *rplD* gene (Gly49His, Lys65Ala) |
| rplD_G68H+K65A_R | CGCGGCCCGTATGCTTCTGCGCGTACATCTTGGCG | |
| phzE_pSRK_GA_F | ATAACAATTTCACACAGGAAACAGCATATGGGTAAATCTGAAAGCG | Molecular cloning of *phzE* gene into the pSRK plasmid (Gibson Assembly) |
| phzE_pSRK_GA_R | CGAGGTCGACGGTATCGATATCATTTGGAAGCAAACG | |

**Supplementary Table 2 | Cryo-EM data collection and refinement statistics.**

| | ***Eco* 70S-PHZ** |
|---|---|
| Microscope | FEI Titan Krios |
| Accelerating Voltage (kV) | 300 |
| Detector | GATAN K2 Summit |
| Spherical aberration (mm) | 2.7 |
| Magnification | 215,000x |
| Defocus range (µm) | -0.6 to -2.0 |
| Micrographs | 2489 |
| Particles picked | 112,130 |
| Particles refined | 65,393 |
| Resolution achieved (Å) | 2.87 |
